# Aggregate data modelling: a fast implementation for fitting pharmacometrics models to summary-level data in R

**DOI:** 10.1101/2025.10.28.685047

**Authors:** Hidde van de Beek, Pyry A.J. Välitalo, J.G. Coen van Hasselt, Laura B. Zwep

**Author notes:** Equal contribution (Hidde van de Beek, Pyry A.J. Välitalo). Telephone: Not given.

## Abstract

Pharmacometric modelling is traditionally performed using individual level data. Recently a new method was developed to fit pharmacometric models to summary level – or aggregate – data. This methodology allows for jointly modelling different data sources, once transformed into aggregate data. As such, the method can be applied to a combination of individual data, pharmacometric models, and aggregate data. In this study we aimed to (1) implement this methodological framework into an accessible R package (admr) and (2) develop a novel algorithm with enhanced computational efficiency. The developed R-package allows calculating aggregate data from different data sources, jointly fitting one or multiple data sources and assessing model performance. The implementation of the newly developed algorithm improves computational efficiency by iteratively reweighting internal Monte Carlo predictions. Three simulation scenarios using different data generating models demonstrated an improvement of 3 to 100-fold speed-up when using the novel Iterative Reweighting Monte Carlo (IR-MC) algorithm, while maintaining the convergence properties of the original MC algorithm. These analyses demonstrated that estimation with the IR-MC algorithm is increasingly more efficient as model complexity rises as compared to the standard MC algorithm, indicating the utility for more complex pharmacometric models. In conclusion, the aggregate data modelling implementation in the admr R package allows for a fast and user-friendly application of the aggregate data modelling framework.

## Introduction

In pharmacometrics, population pharmacokinetic (popPK) modelling usually relies on individual-level data to estimate population parameters and interindividual variability. Since patient-level data are not easily shared, while summary-level data can be, an estimation method was previously proposed enabling pharmacometric models to be fitted to summary-level data and aggregate data [1]. Here, summary-level data consists of summary measures, derived from individual data points, whereas aggregate data refers to summary level data either from a single study or from multiple studies combined. The summary measures used for this modelling technique are the mean and standard deviations of the drug concentration values at specific time points, and the optional addition of the covariance between the concentrations at those timepoints. These covariances are currently not widely available, however, these data can also be calculated from individual patient data or derived from existing pharmacometric models. The latter is particularly valuable since such models are often more readily available than patient-level data. By using literature-reported pharmacometric models, the required mean observations and variability estimates can be generated under the study specific conditions using Monte Carlo (MC) sampling (**Figure 1A**). These estimates can be used in the aggregate data method to combine data from different sources (**Figure 1B**), which allows for pharmacometric analyses, even when direct access to individual-level data is unavailable.

**Figure 1.**
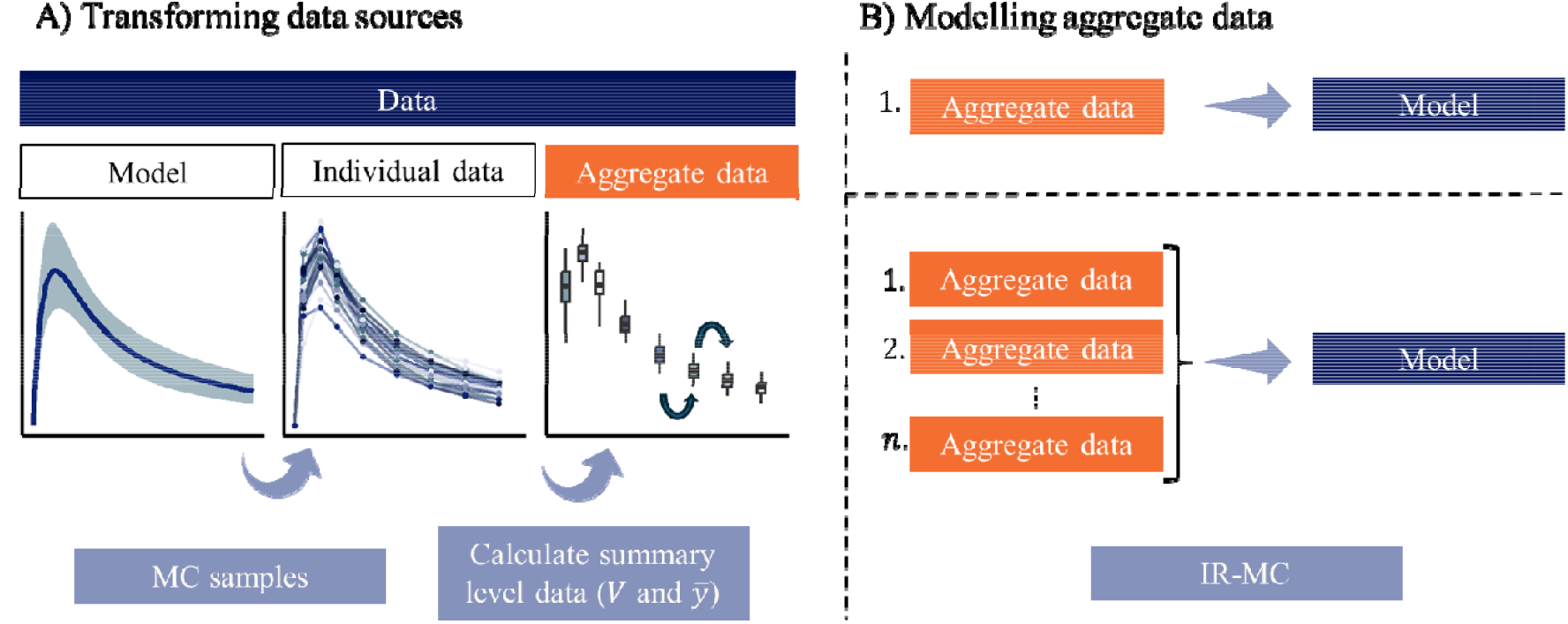
Modelling workflow of (A) transforming different data sources into aggregate data and (B) modelling aggregate data from one or multiple sources. Panel A shows how a model can be used to generate a large number Monte Carlo (MC) samples to reconstruct a population without sampling error. Summary statistics -- (variance-covariance matrix) and (mean vector) -- can then be calculated based on the generated or observed population. Panel B shows the implementation of the developed IR-MC algorithm where one or multiple sources of aggregate can be combined to jointly fit the pharmacometric model.

The previously developed aggregate data MC algorithm is based on loglikelihood estimation and was shown to asymptotically converge to true parameter estimates when the data-generating and analytic models are aligned, providing a robust foundation for handling summary-level data [1]. However, calculation of the loglikelihood for each iteration requires computation of a high number of MC predictions, and computational power can thus become a bottleneck for fitting models to aggregate data. This computational burden limits the method’s scalability, particularly for more complex models or larger datasets. For broader adoption and practical usability, a more computationally efficient algorithm based on reusing the MC predictions is essential.

This study aims to advance the applicability of the aggregate-data modelling framework by: (1) implementing the methodology into an accessible R package (admr) and (2) developing and evaluating an improved algorithm with enhanced computational efficiency.

## Methods

### Package development

To facilitate the implementation of the methodology for aggregate data modelling, a new R package “admr” was developed. This package provides a structured method for fitting pharmacometric models to summary-level data, enabling the estimation of population parameters from aggregate observations. The admr package includes the following functionalities.

- **Data transformation:** Calculating aggregate data from individual-level data or reconstructed from published literature models using Monte Carlo (MC) simulations (**Figure 1A**). These MC simulations can be set to a large number to correct for sampling error in the derived summary data.
- **Model fitting:** Estimation of model parameters from one or multiple data sources (**Figure 1B**). To ensure integration into existing pharmacometric modelling methods, the rxode2 library [2] can be employed for model specification. Models can also be specified using other fast techniques such as C++. An optimized function using the new IR-MC algorithm introduced in this paper is available for extended loglikelihood exploration (Appendix A).
- **Model evaluation**: Evaluation of model performance. Since no individual observations are made, not every standard evaluation metric or visualization method is available. Therefore, other options including tools for assessing goodness-of-fit, parameter convergence, and summary data residuals are available.

The admr package was implemented in R and developed to support broader applications of aggregate data modelling in pharmacometric research. To ensure easy access to the source code and documentation the package is hosted on GitHub. The repository provides detailed installation instructions, usage examples, and model development guidelines (https://github.com/vanhasseltlab/admr). To accompany the development of the R-package, the faster IR-MC algorithm was derived, improving on the MC algorithm used in previous work.

### Algorithm development

An iterative reweighting MC (IR-MC) algorithm was developed for the aggregate data modelling framework. Within this framework, the study design plays a central role in shaping the observed data, since data from subjects sharing the same study design can be aggregated together. A study design represents the independent variables of the dataset, such as sampling schedules or dosing regimens. For *N* individuals, each subject has an elementary design, which may be shared across the population. These designs define the structure of the aggregated responses and their variance-covariance relationships, as summarized by the observed statistics, the means of the concentrations (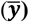), and the variance-covariance matrix (***V***). The IR-MC algorithm estimates population parameters through maximizing the loglikelihood of the estimated summary metrics (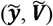) based on *N* individuals sharing the exact same study design (Equation 1).

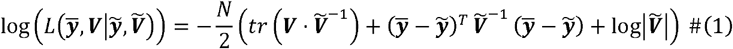

This expression evaluates the goodness-of-fit between observed and predicted aggregate data while accounting for variability and correlations captured in the mean and variance-covariance matrices. The proposed new algorithm contains the following steps to maximize the log-likelihood:

- **Step 1**: MC predictions are computed using the current parameter estimates.
- **Step 2**: While keeping the MC predictions constant, the parameters affect MC predictions through importance weighting. The residual error terms are allowed to directly affect the variance-covariance matrix. These operations are then used to update the predicted mean vector, and the predicted variance-covariance matrix as a function of parameters. Parameters are estimated by maximizing the log-likelihood using the predicted mean vector and variance-covariance matrix.

The MC-algorithm of the original study is a special case where MC samples are not weighted based on relative importance, but all receive equal weights. A detailed theoretical basis of the algorithm improvement is provided in Appendix A, based on the definitions and notations introduced in the original work [1]. Both algorithms are implemented in the R-package as basic functions, alongside an optimized version that employs the IR-MC algorithm for improved exploration in real-world practical applications (Appendix A).

### Simulation study

A simulation study was performed to evaluate the proposed algorithm for three different models. Using the package’s basic implementations of both algorithms, we benchmarked the computational efficiency and convergence of the IR-MC algorithm compared to the original MC algorithm. The simulation scenarios were based on three different data generating models, from which the summary level data were calculated through MC sampling. For data generation and analysis, analytical and ODE models were specified in C++. Each scenario was simulated 20 times. The source code for the simulation study is available as an electronic supplement.

#### Two compartmental model

The first model is an analytical two-compartment model with first-order absorption and elimination. This model was used to simulate patient-level pharmacokinetic (PK) data with fixed effects as shown in **Table 1**. Log-normally distributed IIV with a log-standard deviation was applied to all parameters, with no correlations between the random effects. From this model 5000 individuals were simulated with an administration dose of 100 mg. A dense sampling design (sampling times: 0.1, 0.25, 0.5, 1, 2, 3, 5, 8, and 12 hours) was chosen with precise characterization of the absorption, distribution, and elimination phases.

**Table 1.** The true parameter values of the models used in the three simulation scenarios. The first simulation scenario is a two-compartment model with first-order absorption and elimination, applying random effects on all the fixed effect parameters. The second model has the same structure but without random effects on peripheral volume and intercompartmental clearance. The third simulation scenario is a two-compartment model with a transit compartment absorption process with random effects on all fixed effect parameters.

| Parameter (unit) | Two compartmental model |  | Simplified two compartmental model |  | Transit absorption model |  |
| --- | --- | --- | --- | --- | --- | --- |
|  | Fixed effect | Random effect SD | Fixed effect | Random effect SD | Fixed effect | Random effect SD |
| Absorption rate (L/h) | 1 | 0.3 | 1 | 0.3 | 2 | 0.3 |
| Central volume (L) | 10 | 0.3 | 10 | 0.3 | 10 | 0.3 |
| Peripheral volume (L) | 30 | 0.3 | 30 | - | 30 | 0.3 |
| Intercompartmental clearance (L/h) | 10 | 0.3 | 10 | - | 10 | 0.3 |
| Clearance (L/h) | 5 | 0.3 | 5 | 0.3 | 2 | 0.3 |
| Number of transit compartments |  |  |  |  | 5 | 0.3 |
| Mean transit time |  |  |  |  | 0.5 | 0.3 |
| Proportional error | 10% |  | 10% |  | 10% |  |

#### Simplified two compartmental model

In the second model, the inter-individual variability (IIV) on some fixed effect parameters was set to zero, to enable benchmarking the capability to estimate fixed effects that are not associated with random effects. The same two-compartment model, drug dosing, and sampling design were used, but the IIV was removed from the peripheral volume and inter-compartmental clearance parameters (**Table 1**).

#### Transit absorption model

The third model involved a complex ODE-based absorption model to further investigate the extent of speed-up in a computationally intensive scenario. A two-compartment model with a transit compartment absorption process was used (**Table 1**). Log-normally distributed IIV with a log-standard deviation was applied to all parameters, with no correlations between random effects. In order to ensure a computationally intensive scenario, a steady state dosing and sampling design were used. The sampling times were set between 168 and 176 hours in 0.2-hour intervals, followed by additional sampling between 177 and 190 hours in 1-hour intervals. From this model, 5000 individuals were simulated with an administration dose of 100 mg each 8 hours.

### Evaluation

To compare the newly developed IR-MC algorithm with the previous MC approach, we evaluated parameter convergence, visual predictive checks (VPCs), and computational time.

Convergence was evaluated based on whether the algorithms consistently reached stable and accurate parameter estimates across all runs. Estimation accuracy was quantified using the relative bias (%) of the final predicted estimates versus the true parameters.

VPCs were created for the IR-MC algorithm to assess whether the dynamics of the model were appropriately captured by the aggregate data modelling framework. The data generating models were used to generate a benchmark population plot for the predicted models by the IR-MC algorithm. In the resulting population plots it can be assessed whether the dynamics of each data generating model were captured by the aggregate data modelling framework.

Computational efficiency was quantified by comparing total run time of both algorithms per simulation. The same hardware was used for each model scenario and algorithm, ensuring that differences in execution speed were attributable to algorithmic improvements rather than variations in computational resources.

Overall performance per scenario was assessed across 20 runs, with each run using different initial values generated by adding normally distributed noise (mean = 0, SD = 0.1) to the log-transformed parameter values. Visualization and data wrangling was performed using the tidyverse package [3].

## Results

### Parameter estimation

In order to assess the convergence properties, we assess the relative bias in both algorithms. With initial values that were equal to the true parameter values, both the IR-MC and MC algorithms successfully converged to the parameter values. When initial values were changed, in all three simulation scenarios the estimated parameters remained consistent across different initial values, only showing a very slight bias of below 1% for the absorption rate, absorption rate omega and the proportional error (**Figure 2**). A similar trend was observed for the other parameters (**Figure A1, Figure A2, Figure A3**). Both algorithms performed similarly. This result highlights the robustness of the algorithms in handling different model structures.

**Figure 2.**
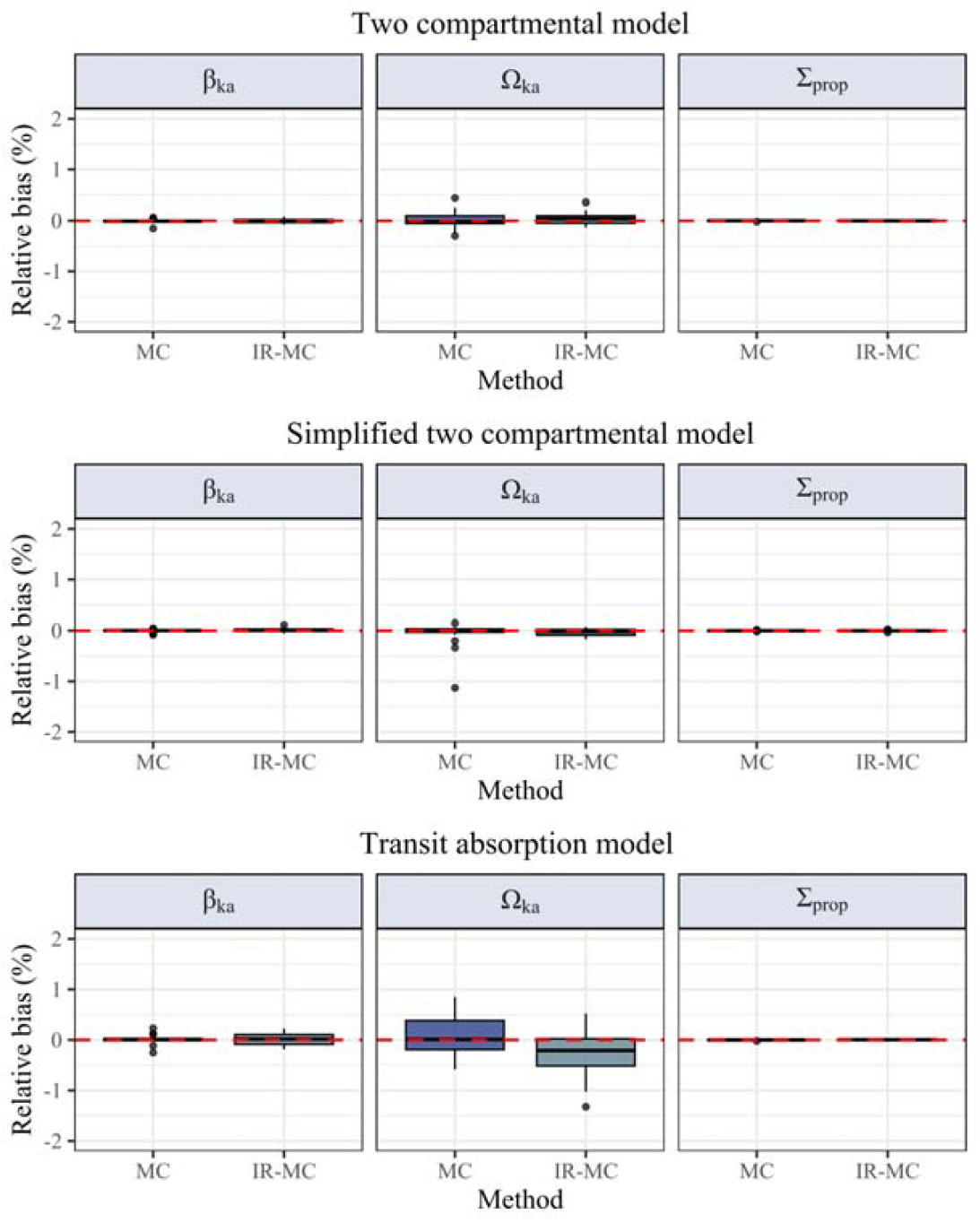
Relative bias for the three simulation scenarios on estimates of absorption rate, associated random effect, and the proportional error of the model for 20 simulations, estimated using two methods (MC, IR-MC). Bias is calculated as a percentage of the true parameter. Each model shows a boxplot of near-zero bias in the fixed effect and random effect.

In order to evaluate how accurately the model captures the underlying dynamics, the population pharmacokinetic profiles for the true models and the fitted models were compared (**Figure 3**). These VPCs of the IR-MC predictions for the three simulation scenarios show identical model predictions over all iterations compared to the true model. There is no clear indication of model misspecification. These results suggest that the small systematic bias in parameter estimates does not lead to bias in model predictions.

**Figure 3.**
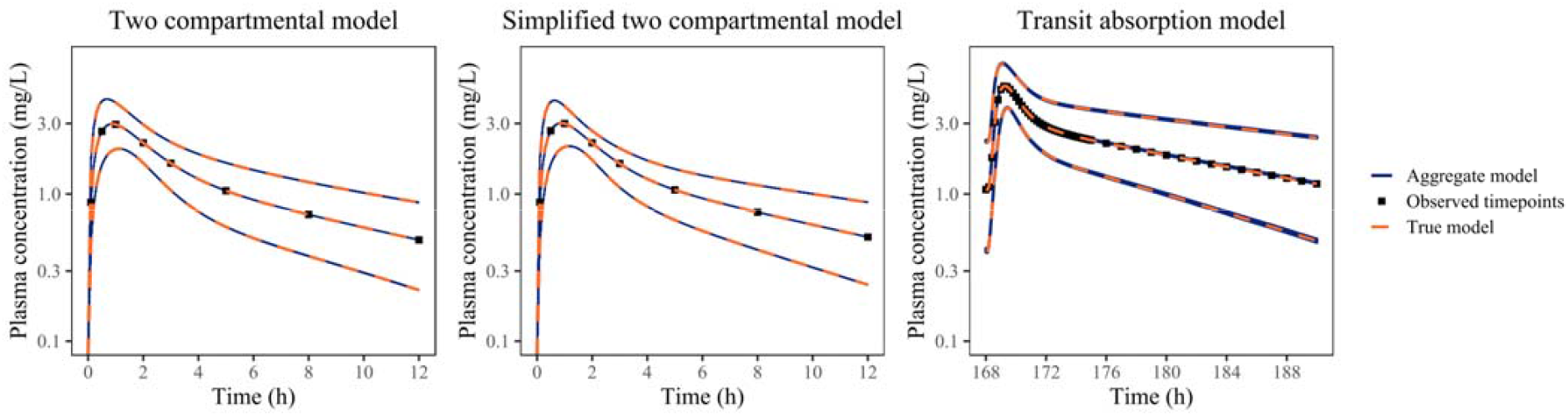
Pharmacokinetic profiles of the true model and aggregate data-based model estimated from the IR-MC algorithm. The panels show the 5th, 50th, and 95th percentiles of the true model and estimated model for each simulation scenario.

### Computational Efficiency

In order to assess computational efficiency of the new IR-MC algorithm compared to the previous MC algorithm, we calculated the computation time and time differences for each simulation study. In all models the newly developed IR-MC algorithm outperformed the previous MC algorithm (**Table 2**). In the two compartmental model (Simulation 1) the IR-MC algorithm achieved an average 9-fold reduction in time to convergence compared to the MC algorithm. When removing two IIV parameters (Scenario 2), the time to convergence reduces to a 4-fold change. The transit absorption model (Scenario 3) showed the computational performance of the IR-MC algorithm when applied to a more complex pharmacokinetic model using ODEs, resulting in a 144-fold reduction in time to convergence. These results show an increase in computational efficiency with an increase in model complexity. The ability of the IR-MC algorithm to handle transit compartment models efficiently suggests its applicability to a broad range of pharmacokinetic modelling scenarios requiring computationally intensive solutions.

**Table 2.**
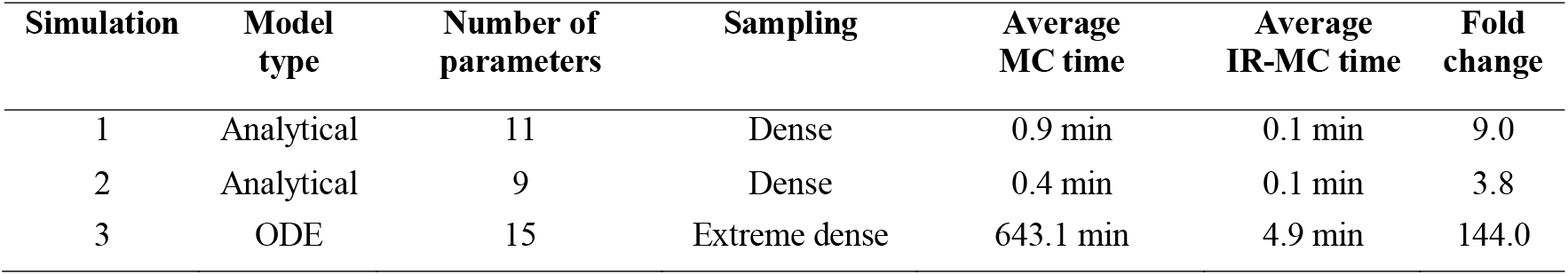
Computation times and time differences of the IR-MC algorithm and the MC algorithm for the three simulation scenarios. The number of parameters reflects model complexity.

| Simulation | Model type | Number of parameters | Sampling | Average MC time | Average IR-MC time | Fold change |
| --- | --- | --- | --- | --- | --- | --- |
| 1 | Analytical | 11 | Dense | 0.9 min | 0.1 min | 9.0 |
| 2 | Analytical | 9 | Dense | 0.4 min | 0.1 min | 3.8 |
| 3 | ODE | 15 | Extreme dense | 643.1 min | 4.9 min | 144.0 |

## Discussion

In the current study, we reported on the development of a new R-package, admr, to facilitate aggregate modelling framework for compartmental population PK/PD models, enabled by a previously proposed quantitative methodology [1]. The admr R package allows for data handling, estimation of compartmental population models, and visual evaluation of model fit. The usage of the rxode2 library for model specification ensures the integration of the existing pharmacometric modelling ecosystem. The IR-MC algorithm implemented for model fitting implemented in the current study improved efficiency for practical application of the method.

The package includes visual evaluation tools, which can be used to check the goodness-of-fit. Since individual-level data are not applicable within this framework, standard residual-based diagnostic plots (e.g., individual residuals vs. time or predictions) cannot be performed. Consequently, model evaluation relies primarily on alternative diagnostics, including: (1) comparison of observed and predicted confidence intervals, (2) convergence diagnostics across multiple chains, and (3) graphical comparisons of observed versus predicted means and covariance structures. The summary residual diagnostics, such as the mean residual at each time point and the residual variance– covariance matrix, may be used to assess overall goodness-of-fit. Further work can be done to improve the development and interpretation of these model diagnostics.

The IR-MC algorithm was able to accurately estimate the model parameters across all three simulation scenarios. Notably, it achieved performance equivalent to the original MC algorithm, but with improved computational efficiency. This efficiency gain is a direct effect of being able to re-use the existing MC predictions with updated weightings, instead computing a new set of MC predictions at each parameter update [4]. The results of the three scenarios collectively show that the relative speedup increases with model complexity. Furthermore, the three simulation scenarios, differing in modelling and computational complexity, show us a range of possible applications of the framework and IR-MC algorithm. This study aimed to compare two different estimation algorithms on performance, therefore, we empirically evaluated a small set of possible popPK models. The performance is expected to generalize to other popPK models, but this has not been evaluated thoroughly.

Future extensions of this framework could significantly improve the utility of aggregate data modelling in complex pharmacometric analyses. On such extension is in the domain of covariate modelling, as most population PK/PD analyses seek to identify patient covariates which can explain inter-individual variability observed for specific PK or PK/PD model parameters [5] to inform dosing strategies [6]. Aggregate covariate modelling is not yet incorporated in the current theoretical framework but is under active development. Additionally, the inclusion of, for example, inter-occasion variability within the modelling framework would allow for more accurate representation of variability.

In conclusion, we introduced a fast method for fitting pharmacometric models to (a combination of) summary-level data, population models and individual level data. The open-source R package implementation ensures accessibility, allowing for future broader adoption within the pharmacometric community. The ability to integrate different data and models as input makes for a particularly valuable new tool for meta-analyses and model-based drug development.

## Supporting information

electronic supplement

## Appendix A

This Appendix outlines the detailed theoretical basis for the IR-MC algorithm.

### Iterative reweighting MC algorithm for aggregate data

The algorithm estimates population parameters through maximizing the logli keliho od of the estimated summary metrics 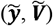 based on *N* individuals sharing a study design. In this method 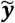 and 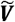 are calculated as a Monte Carlo integral of elementary predictions 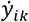 weighted by a vector ***w***

Here, 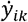 represents the prediction for the k-th measurement or output based on the i-th Monte Carlo sample of the individual parameter vector, computed as:

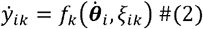

With the individual parameter vector 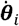 is defined as:

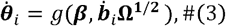

where *g* (.) is a function defining the individual parameters as a function of fixed effects and random effects, **Ω**^**½**^ is the Cholesky decomposition of **Ω** and 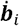 is the ith set of Sobol-sampled vectors with a mean of zero and standard deviation of one [7, 8]. We use **Ψ** to define the set of all parameters, including the fixed-effects vector ***β***, random effects variance-covariance matrix ***Ω*** and the residual variance matrix **∑**.

Using the elementary predictions 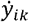 calculated in equation 2, the predicted mean at the ***K*** -th measurement is computed by:

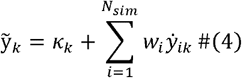

Equation 4 introduces the term *k*_*k*_, which is explained in detail in equation 8. For now, we briefly mention that this term is necessary for the estimation of fixed effects parameters that are not associated with a random effect.

The predicted variance-covariance matrix 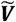 can be computed element by element. The covariance between ***k***-th ***l***-th measurement, 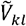 can be calculated as

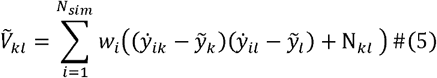

Where N_*kl*_ is the impact of the residual variability on the diagonals of 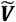 and is calculated by linearizing the residual error function h(.) with respect to the residual error term as follows:.

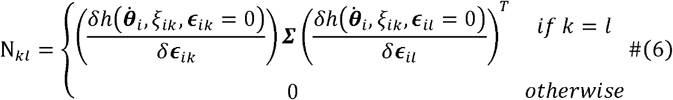

The weights *W*_*i*_ of the previously generated elementary predictions can be updated based on current parameter estimates **Ψ** Each weight *W*_*i*_ reflects the relative importance of sample 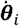 by comparing the likelihood of the sample under the target distribution 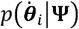 with its likelihood under the original proposal distribution 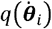:

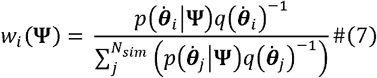

Thus, it is possible to generate an updated set of predictions on the basis of previously generated elementary predictions calculated on the basis of previously generated individual parameters 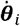 and on the basis of a new set of parameters **Ψ** by emphasizing those individual parameter values that are more likely under the new set of **Ψ** when compared to the parameters that were used in obtaining the previously generated elementary predictions. This allows the reweighting of all individual parameters associated with a random effect.

In the case wherein an individual parameter is not associated with a corresponding random effect, in our opinion the most efficient solution is to compute a correction term ***k*** to account for fixed effects that apply consistently across all individuals, without inter-individual or inter-occasion variability. Elements of ***k*** were already included in equation 4. Now we outline how the correction factor for the k-th prediction is calculated.

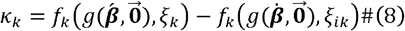

Where 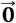 is a vector of zeros with length equal to the number of random effects, 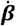 is the previous set of fixed effects used to generate the set of elementary predictions, and 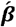 is a vector consisting of a mix of both previous and current fixed-effects estimates; the fixed effects not associated with random effects are set to current estimates, whereas the rest of the fixed effects are set to previous values that were used to generate the set of elementary predictions. This *k*_*k*_ only reflects the differences in predictions between the previous and the updated fixed-effects parameters not associated with random effects. The value of *k*_*k*_ is fast to compute, because it is not integrated over the random effects values but rather computed with an assumption of random effects being zero.

The eventual IR-MC algorithm updates parameters in two main steps, which are iterated through until a convergence criterion is met:

- **Step 1**: Elementary predictions 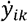 are computed using the current parameter estimates according to equations 2 and 3, with an option to expand the current values of **Ω** by a scaling factor *ρ* which broadens the proposal distribution, allowing the iterative reweighting algorithm to explore beyond the strict proposal space implied by the current parameter estimates, and to accelerate convergence. The probability densities of the proposal distribution are stored into memory and treated as constant.
- **Step 2**: Treating the elementary predictions 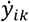 and proposal distribution densities as constant, the parameters **Ψ** affect predictions through weights which are calculated according to equation 7, through correction factor ***k*** as defined in equation 8, and through residual error contribution terms as defined in equation 6. These terms are then used to update the predicted mean vector according to equation 4, and the predicted variance-covariance matrix according to equation 5. Parameters are estimated by maximizing the log-likelihood according to expression 3 using the predicted mean vector and variance-covariance matrix.

### Original MC algorithm for aggregate data

For the purposes of the aggregate data MC approximation in the original paper, are ***w*** set to the number of MC Samples 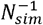, making it a special case of the IR-MC algorithm. When weights are calculated as 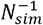, the equations 5 and 6 together correspond to equation 18 from the previous study[1].

Thus, given that 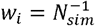 for all of *w*_*i*_ and thus not needing to be updated during step 1, and given that *K*_*k*_ in equation 4 is zero in the case of aggregate data MC approximation, fitting models to data using the aggregate data MC approximation can be accomplished by iterat ively u pdating all the elementary predictions 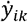 as a function of the current estimates of **Ψ** and then recomputing 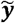 and 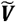 the on the basis of these elementary predictions. Therefore, fitting models to data under maximum likelihood estimation, represents the following optimization problem:

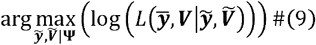

### Optimized IR-MC function for practical application

An optimized function using the new IR-MC algorithm is included in the package. The core algorithm as outlined in the sections above remains the same. However, to ensure proper loglikelihood landscape exploration, two computational steps were added. First, sequential phases using different adaptive search bounds can be specified to more efficiently explore the loglikelihood space. This phased approach helps to map high-probability regions. The time spent in each phase, and the search bounds can be specified by the user. Secondly, multiple independent chains can be initialized with user-defined perturbations (using additive proportional Gaussian noise) applied to the starting values. This implementation increases the searched likelihood space, which enhances likelihood maximization.

## Appendix B

### Two compartmental model

**Figure A1.**
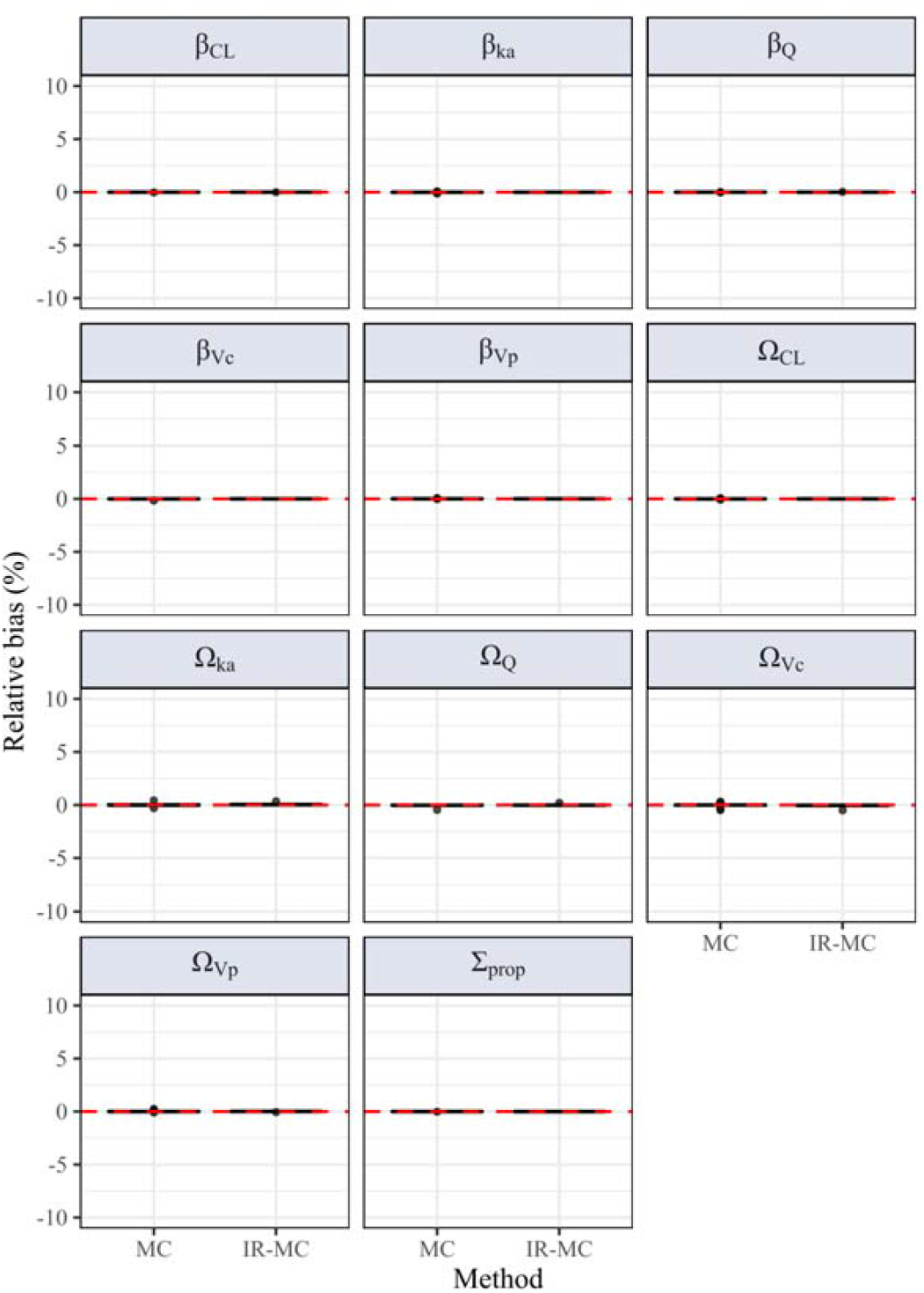
Relative bias for the estimated parameters of the two compartmental model on all fixed and random effects of the model for 20 simulations, estimated using two methods (MC, IR-MC). Bias is measured as a percentage of the true parameter. Each model shows a boxplot of near-zero bias in the fixed effect and random effect.

### Simplified two compartmental model

**Figure A1.**
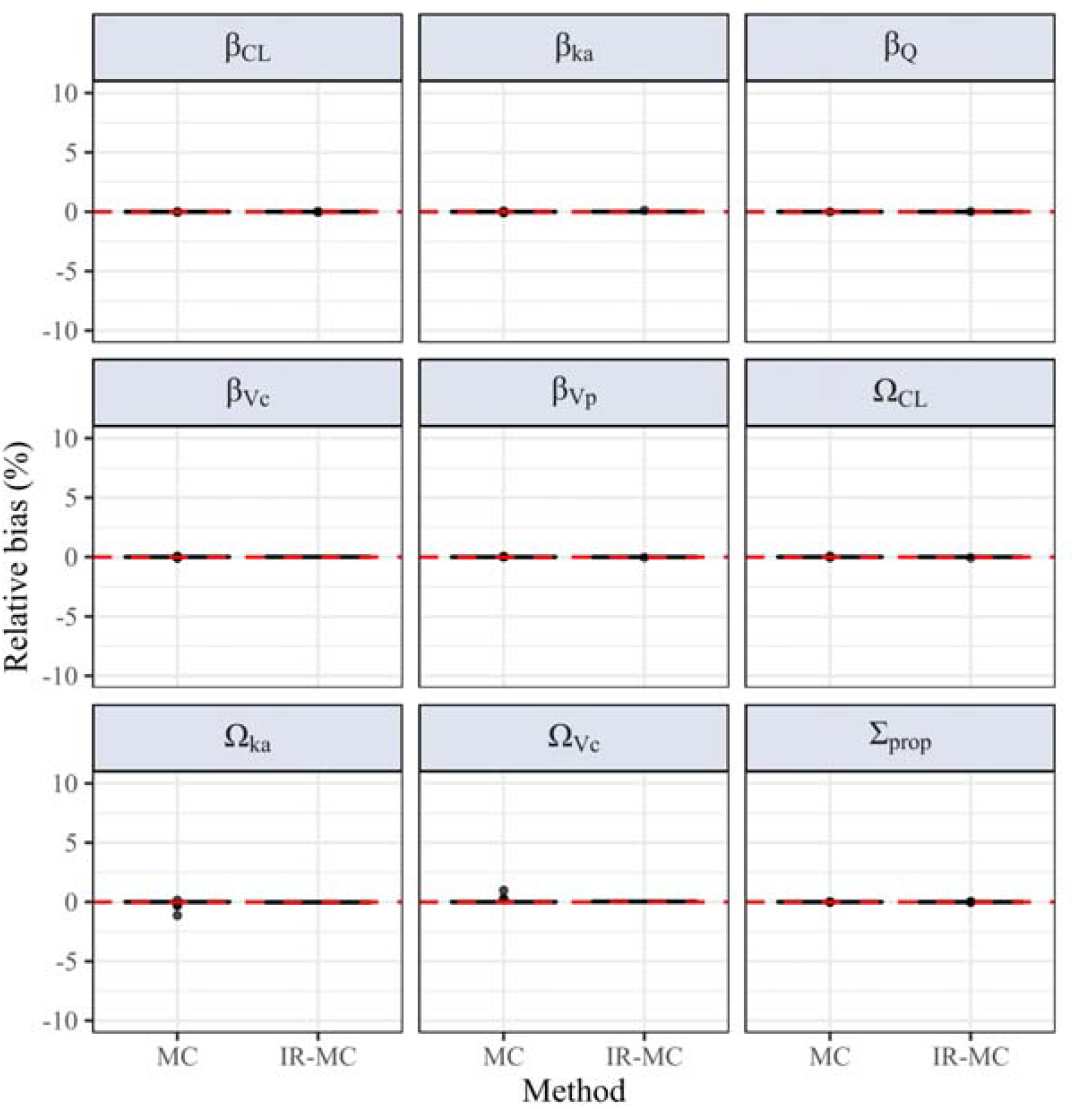
Relative bias for the estimated parameters of the simplified two compartmental model on all fixed and random effects of the model for 20 simulations, estimated using two methods (MC, IR-MC). Bias is measured as a percentage of the true parameter. Each model shows a boxplot of near-zero bias in the fixed effect and random effect.

### Transit absorption model

**Figure A2.**
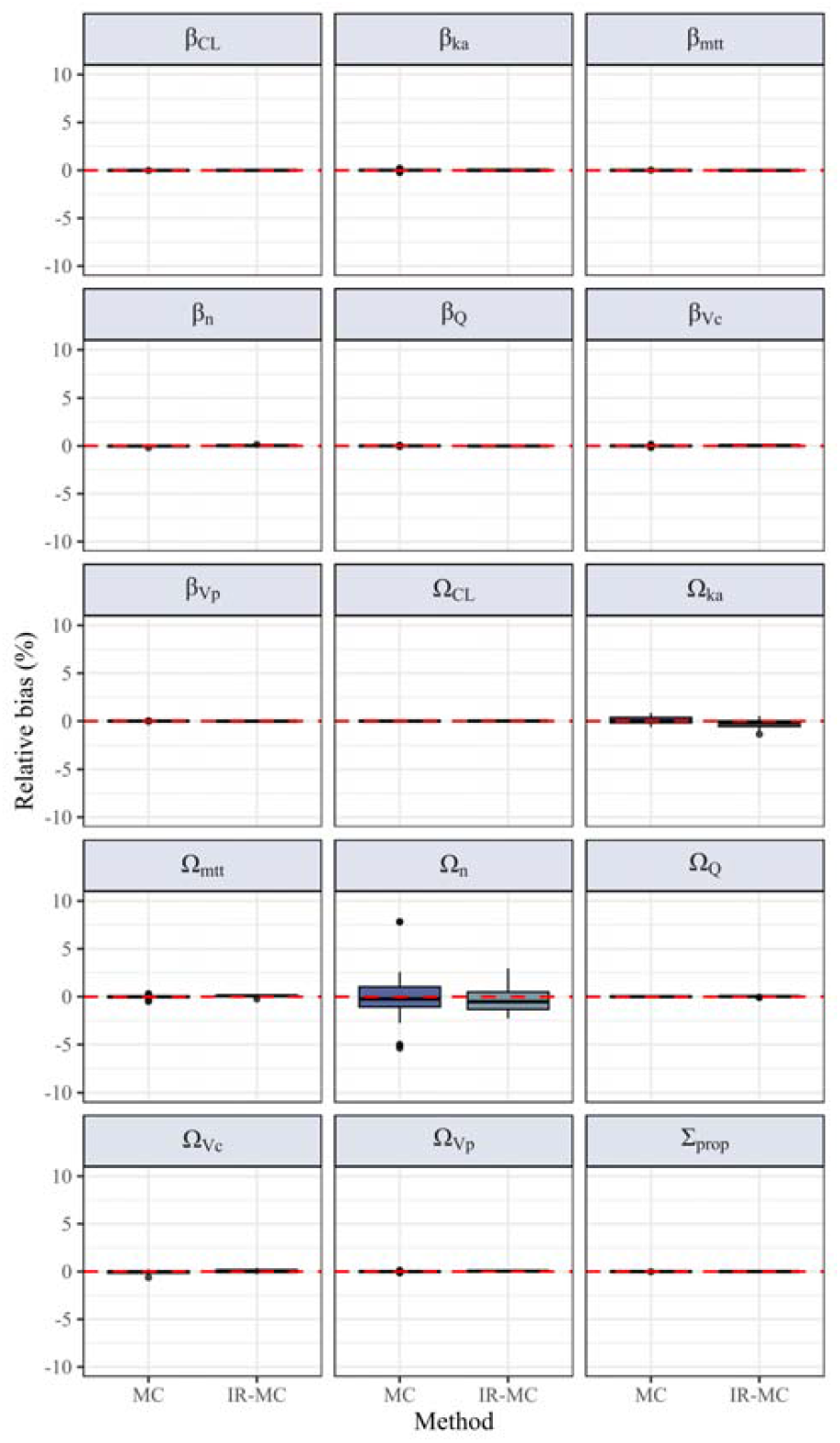
Relative bias for the estimated parameters of the transit absorption model on all fixed and random effects of the model for 20 simulations, estimated using two methods (MC, IR-MC). Bias is measured as a percentage of the true parameter. Each model shows a boxplot of near-zero bias in the fixed effect and random effect.

## References

1. Välitalo PAJ (2021) Pharmacometric estimation methods for aggregate data, including data simulated from other pharmacometric models. J Pharmacokinet Pharmacodyn 48:623–638. 10.1007/s10928-021-09760-1

2. Fidler ML, Wang W, Hindmarsh A, et al (2025) rxode2: Facilities for Simulating from ODE-Based Models

3. Wickham H, Averick M, Bryan J, et al (2019) Welcome to the Tidyverse. J Open Source Softw 4:1686. 10.21105/joss.01686

4. Levine RA, Casella G (2001) Implementations of the Monte Carlo EM Algorithm. J Comput Graph Stat 10:422–439. 10.1198/106186001317115045

5. Joerger M (2012) Covariate Pharmacokinetic Model Building in Oncology and its Potential Clinical Relevance. AAPS J 14:119–132. 10.1208/s12248-012-9320-2

6. Wade JR, Beal SL, Sambol NC (1994) Interaction between structural, statistical, and covariate models in population pharmacokinetic analysis. J Pharmacokinet Biopharm 22:165–177. 10.1007/BF02353542

7. Sobol IM (1976) Uniformly distributed sequences with an additional uniform property. USSR Comput Math Math Phys 16:236–242. 10.1016/0041-5553(76)90154-3

8. Renardy M, Joslyn LR, Millar JA, Kirschner DE (2021) To Sobol or not to Sobol? The effects of sampling schemes in systems biology applications. Math Biosci 337:108593. 10.1016/j.mbs.2021.108593

