## Supplementary material for "Aggregate data modelling: a fast implementation for fitting pharmacometrics models to summary-level data in R": electronic supplement: cpp.R

```
cppFunction('NumericVector Cpp_twocomp(NumericVector pars,NumericVector ti,bool ss) {
  NumericVector out(ti.size());
  double cl=pars[0];double v1=pars[1];double v2=pars[2];double q=pars[3];double ka=pars[4];
  double k=cl/v1;
  double k12=q/v1;
  double k21=q/v2;
  
  double beta = 0.5*(k+k12+k21-sqrt((k+k12+k21)*(k+k12+k21)-4*k*k21));
  double alpha = k*k21/beta;
  
  double A = (alpha-k21)/(alpha-beta)/(ka-alpha)*ka/v1;
  double B = (beta-k21)/(beta-alpha)/(ka-beta)*ka/v1;
  
  for (int i = 0; i < ti.size(); ++i) {
    
    out[i]=A*exp(-ti[i]*alpha)+B*exp(-ti[i]*beta)-(A+B)*exp(-ka*ti[i]);
    
    if (ss) {
      out[i]= (A*exp(-24*alpha)+B*exp(-24*beta)-(A+B)*exp(-ka*24))/(1-exp(-24*beta));
    }
  }
  return out;
}')

cppFunction('
#include <Rcpp.h>
#include <cmath>
#include <vector>
#include <omp.h>  // OpenMP for parallelization
using namespace Rcpp;

// [[Rcpp::plugins(openmp)]]
// [[Rcpp::export]]
NumericMatrix Cpp_twocomp2(NumericVector cl, NumericVector v1, NumericVector v2,
                               NumericVector q, NumericVector ka, NumericVector ti, bool ss) {
  int n_ind = cl.size();  // Number of individuals
  int n_time = ti.size(); // Number of time points
  NumericMatrix out(n_ind, n_time);

  #pragma omp parallel for  // Parallelize across individuals
  for (int iter = 0; iter < n_ind; ++iter) {
    double k = cl[iter] / v1[iter];
    double k12 = q[iter] / v1[iter];
    double k21 = q[iter] / v2[iter];

    double sum_rates = k + k12 + k21;
    double sqrt_disc = sqrt(sum_rates * sum_rates - 4 * k * k21);
    double beta = 0.5 * (sum_rates - sqrt_disc);
    double alpha = k * k21 / beta;

    double A = (alpha - k21) / ((alpha - beta) * (ka[iter] - alpha)) * ka[iter] / v1[iter];
    double B = (beta - k21) / ((beta - alpha) * (ka[iter] - beta)) * ka[iter] / v1[iter];

    double ss_factor = 1.0 / (1 - exp(-24 * beta)); // Precompute steady-state correction factor

    for (int i = 0; i < n_time; ++i) {
      double exp_alpha = exp(-ti[i] * alpha);
      double exp_beta  = exp(-ti[i] * beta);
      double exp_ka    = exp(-ka[iter] * ti[i]);

      out(iter, i) = A * exp_alpha + B * exp_beta - (A + B) * exp_ka;

      if (ss) {
        double exp_24_alpha = exp(-24 * alpha);
        double exp_24_beta  = exp(-24 * beta);
        double exp_24_ka    = exp(-ka[iter] * 24);

        out(iter, i) = (A * exp_24_alpha + B * exp_24_beta - (A + B) * exp_24_ka) * ss_factor;
      }
    }
  }

  return out;
}
')

cppFunction('
NumericVector Cpp_transit_dt(NumericVector pars, NumericVector ti) {
  int n_out = ti.size();
  NumericVector out(n_out);
  
  // Extract parameters
  double cl = pars[0], vc = pars[1], vp = pars[2], q = pars[3], ka = pars[4];
  double mtt = pars[5], n = pars[6];
  
  // Compute rate constants
  double k = cl / vc;
  double k23 = q / vc;
  double k32 = q / vp;
  double ktr = (n + 1) / mtt;   // rate for gamma PDF
  double bio = 1.0;
  double podo = 100.0;  // Dose amount
  
  // Initialize compartments: depot, central, peripheral
  double depot = 0.0, cen = 0.0, periph = 0.0;
  
  // Use a small time step for integration
  double dt = 0.025;
  
  // Container for dosing times
  std::vector<double> doseTimes;
  
  // Administer the initial dose at t=0
  doseTimes.push_back(0.0);
  
  // Simulation time starts at 0
  double t = 0.0;
  int j = 0;  // index for output time points
  
  // Run simulation until we pass the last requested time
  while (t <= ti[n_out - 1] + dt) {
    // Every 24 hours (after t=0) add a new dose event.
    if (t > 0.0 && fmod(t, 24.0) < dt) {
      doseTimes.push_back(t);
    }
    
    // Sum contributions from all doses in the transit chain.
    // For a dose given at time t_dose, the contribution at time t is:
    //   bio * podo * ktr * (ktr*(t - t_dose))^n * exp(-ktr*(t - t_dose)) / gamma(n+1)
    double depot_input = 0.0;
    for (size_t i = 0; i < doseTimes.size(); i++) {
      double age = t - doseTimes[i];
      if (age >= 0) {
        depot_input += bio * podo * ktr * pow(ktr * age, n) * exp(-ktr * age) / exp(lgamma(n + 1));
      }
    }
    
    // Update depot using Eulers method:
    depot += depot_input * dt - ka * depot * dt;
    depot = std::max(depot, 0.0);  // avoid negative values
    
    // Update central and peripheral compartments:
    double dCen = ka * depot - (k + k23) * cen + k32 * periph;
    double dPeriph = k23 * cen - k32 * periph;
    cen += dCen * dt;
    periph += dPeriph * dt;
    
    // Advance time by dt
    t += dt;
    
    // When we cross an output time, store the plasma concentration
    while (j < n_out && t >= ti[j]) {
      out[j] = cen / vc;
      j++;
    }
  }
  
  return out;
}
')

cppFunction('
#include <Rcpp.h>
#include <cmath>
#include <vector>
#include <omp.h>

using namespace Rcpp;
using namespace std;

// [[Rcpp::plugins(openmp)]]

// [[Rcpp::export]]
NumericMatrix Cpp_transit2_dt(NumericVector cl, NumericVector vc, NumericVector vp, NumericVector q, 
                           NumericVector ka, NumericVector mtt, NumericVector n, NumericVector ti) {
  int n_ind = cl.size();  // Number of individuals
  int n_time = ti.size(); // Number of time points
  omp_set_num_threads(10);
  NumericMatrix out(n_ind, n_time); // Output matrix

  double dt = 0.025;  // Small fixed time step for accuracy

  // Parallel processing of individuals using OpenMP
  #pragma omp parallel for
  for (int i = 0; i < n_ind; i++) {
    // Compute individual-specific parameters
    double k = cl[i] / vc[i];
    double k23 = q[i] / vc[i];
    double k32 = q[i] / vp[i];
    double ktr = (n[i] + 1) / mtt[i];
    double bio = 1.0;
    double podo = 100.0;  // Dose amount

    // Initialize compartments
    double depot = 0.0, cen = 0.0, periph = 0.0;

    // Container for dosing times (thread-safe)
    vector<double> doseTimes;
    doseTimes.push_back(0.0); // Initial dose at t=0

    // Time variable
    double t = 0.0;
    int j = 0;  // Index for output time points

    // Loop until all requested times are covered
    while (t <= ti[n_time - 1] + dt) {
      // Every 24 hours, add a new dose event
      if (t > 0.0 && fmod(t, 24.0) < dt) {
        doseTimes.push_back(t);
      }

      // Compute depot input from all past doses
      double depot_input = 0.0;
      for (size_t d = 0; d < doseTimes.size(); d++) {
        double age = t - doseTimes[d];
        if (age >= 0) {
          depot_input += bio * podo * ktr * pow(ktr * age, n[i]) * exp(-ktr * age) / exp(lgamma(n[i] + 1));
        }
      }

      // Update depot using Eulers method
      depot += depot_input * dt - ka[i] * depot * dt;
      depot = max(depot, 0.0);  // Avoid negative values

      // Update central and peripheral compartments
      double dCen = ka[i] * depot - (k + k23) * cen + k32 * periph;
      double dPeriph = k23 * cen - k32 * periph;
      cen += dCen * dt;
      periph += dPeriph * dt;

      // Advance time
      t += dt;

      // Store output concentrations at specified times
      while (j < n_time && t >= ti[j]) {
        out(i, j) = cen / vc[i];
        j++;
      }
    }
  }

  return out;
}
')
```
