## Supplementary material for "Aggregate data modelling: a fast implementation for fitting pharmacometrics models to summary-level data in R": electronic supplement: internal_functions.R

```
anticovlogit <- function(x) { # Performs logit and inverse logit transformations for values constrained to the [-1,1] domain.
  2*exp(x)/(exp(x)+1)-1
}

backtransfunchooser <- function(fs) { # Returns appropriate back-transformation or derivative functions based on a list of transformation function.
  translist <- c(log=exp,
                 covlogit=anticovlogit,
                 logit=function(x) exp(x)/(1+exp(x)),
                 identity=identity)
  res <- translist[fs]
  for (i in which(map_lgl(res,is.null))) {
    names(res)[i]  <- fs[i]
    res[[i]] <- numtransfun(fs[i])
  }
  res
}

covlogit <- function(x) { # Performs logit and inverse logit transformations for values constrained to the [-1,1] domain.
  x2 <- (x+1)/2
  log(x2/(1-x2))
}

derivtransfunchooser <- function(fs) { # Returns appropriate back-transformation or derivative functions based on a list of transformation names.
  translist <- c(log=exp,
                 covlogit=function(x) 2*exp(x)/(1+exp(x))^2,
                 logit=function(x) exp(x)/(1+exp(x))^2,
                 identity=function(x) 1)
  res <- translist[fs]
  for (i in which(map_lgl(res,is.null))) {
    names(res)[i]  <- fs[i]
    res[[i]] <- function(x) grad(fs[i],x)
  }
  res
}

foceapprEV_single <- function(opts,bi,biseq # Uses the First-Order Conditional Estimation (FOCE) method to approximate expected values for a single individual.
) {
  if (missing(bi)) bi <- matrix(0,1,ncol(opts$p$Omega))
  theta_i <- g_iter(opts,bi)
  Eind <- opts$f(opts$time,theta_i)
  Etyp <- opts$f(opts$time,opts$p$beta)
  d_f_d_bi <- jacobiann(function(et) opts$f(opts$time,g_iter(opts,et)),bi)
  E <- t(c(Eind)-d_f_d_bi %*% t(bi))
  V <- d_f_d_bi %*% opts$p$Omega %*% t(d_f_d_bi)
  if (opts$interact) V <- opts$h(list(V=V,E=Eind),opts$p)$V else V <- opts$h(list(V=V,E=Etyp),opts$p)$V
  res <- list(E=E,V=V,Etyp=Etyp,Eind=Eind,d_f_d_bi=d_f_d_bi,weights=weights)
  res
}

g_iter <- function(opts,bi) { # Applies the model g to generate individual-specific parameter values theta_i based on random effects bi.
  if (!is.matrix(bi)) {
    opts$g(opts$p$beta,bi,opts$ai)
  } else {
    t(apply(bi,1,function(i) opts$g(opts$p$beta,i,opts$ai)))
  }
}

gen_bi <- function(opts,expanded=TRUE) { # Generates random effects bi based on the covariance matrix in the options, with optional expansion.
  Omega <- opts$p$Omega
  if (expanded) Omega <- Omega*opts$omega_expansion
  gen_bi2(Omega,opts$biseq)
}

gen_bi2 <- function(Omega,biseq) { # Generates random effects bi based on the covariance matrix in the options, with optional expansion.
  ## error handling for the  special case of zero omegas
  if (any(diag(Omega)==0))
    diag(Omega)[diag(Omega)==0] <- 1e-10
  biseq %*% chol(Omega)
}

grad <- function(f,var) gradient(f,unname(var)) # Gradient of a function without names.

jacobiann <- function(f,var) jacobian(f,unname(var))

list_to_mat <- function(l) { # Converts a list of values into a matrix by organizing them into a block-diagonal format.
  ## transform a list of values to a band matrix, similar to a series of
  ## NONMEM $OMEGA blocks
  mdim <- map_dbl(l,function(i) 0.5*(sqrt(1+8*length(i))-1))
  m <- matrix(0,sum(mdim),sum(mdim))
  cums <- cumsum(c(0,head(mdim,-1)))
  for (i in seq_along(mdim)) {
    indices <- cums[i]+seq_len(mdim[i])
    mtemp <- matrix(0,mdim[i],mdim[i])
    mtemp[upper.tri(mtemp,diag=TRUE)] <- l[[i]]
    m[indices,indices] <- uptolow(mtemp)
  }
  m
}

logdens2wt <- function(x) { # Converts log-densities into normalized weights for importance sampling.
  x2 <- exp(x-max(x)) ## standardize to avoid floating point errors
  x2[!is.finite(x2)] <- max(x2[is.finite(x2)])
  x2/sum(x2)
}

lowtoup <- function(m) { # Copies the upper triangular part of a matrix to the lower triangular
  m[upper.tri(m)] <- t(m)[upper.tri(m)]
  m
}

mat_to_list <- function(x) { # Converts a matrix back into a list, extracting diagonal and off-diagonal blocks.
  ## transform a sparse matrix to a list of values, similar to list_to_mat
  if (nrow(x)!=ncol(x)) stop("Need square matrix!")
  if (nrow(x)==1) return(list(c(x)))
  check <- apply(x,2,function(i) sum(rle(i!=0)$values)>1)
  if (any(check)) stop("Matrix needs to be either full, band or diagonal; all other types are unsupported!")
  i <- 1
  res <- list()
  while(i<=ncol(x)) {
    l <- sum(x[,i]!=0)
    if (l==0)  l <- 1 ## for the special case of zero omega
    ind <- i-1+seq_len(l)
    newcomp <- c(x[ind,ind][upper.tri(x[ind,ind],diag=TRUE)])
    res <- c(res,list(newcomp))
    i <- i+l
  }
  res
}

maxfunc <- function(opts) {
  bi <- gen_bi(opts)
  origbeta <- opts$p$beta
  rawpreds <- opts$f(opts$time,g_iter(opts,bi))
  NAfilters <- apply(rawpreds,1,function(i) !any(is.na(i)))
  adjust <- !any(is.na(opts$single_betas))
  propdens <- mnorm::dmnorm(bi,mean=rep(0,nrow(opts$p$Omega)),sigma=opts$p$Omega*opts$omega_expansion,log=TRUE)$den

  function(pnew,returnEV=FALSE) {
    pneww <- opts$ptrans(pnew)
    newdens <- opts$p_thetai(pneww,origbeta,bi)
    wttot <- logdens2wt(newdens[NAfilters]-propdens[NAfilters])
    EVnow <- with(cov.wt(rawpreds[NAfilters,],wttot,method="ML"),list(E=center,V=cov))
    EVnow <- opts$h(EVnow,pneww)
    if (adjust) {
      kappa <- opts$f(opts$time,opts$g(ifelse(opts$single_betas,pneww$beta,origbeta))) - rawpreds[1,]
      EVnow$E <- EVnow$E + kappa
    }
    if (returnEV) EVnow else nllfun(opts$obs,EVnow,n=opts$n)
  }
}

MCapprEV <- function(opts) { # Performs a Monte Carlo approximation to compute the expected values (mean and covariance) of the model outputs based on random effects.
  p <- opts$p
  ## p is normal-scale parameters
  bi <- gen_bi(opts)
  theta_i <- g_iter(opts,bi)
  m <- opts$f(opts$time,theta_i)
  if (opts$omega_expansion==1) {
    r <- meancov(m)
  } else {
    wt <- samplogdensfun(bi,p,opts$omega_expansion)
    r <- meancov(m,logdens2wt(wt))
  }
  opts$h(r,opts$p)
}


nllfun <- function(obsEV,predEV,invpredV,n=1) {   # Computes the negative log-likelihood given observed and predicted expected values and their covariance.
  if (missing(invpredV)) invpredV <- solve(predEV$V)
  resids <- c(obsEV$E-predEV$E) ## force to simple vector
  c(1/2*n*(log(det(predEV$V))+sum(diag(obsEV$V %*% invpredV))+t(resids) %*% invpredV %*% resids))
}

numtransfun <- function(origfun) # Creates a back-transformation function numerically based on an original transformation function.
  ## Numeric back-transformation function, inefficient.
  function(x) optimize(function(xx) (xx-do.call(origfun,list(x)))^2,c(-1000,1000))$minimum
translist_string <- c(log="exp",
                      covlogit="2*exp(x)/(exp(x)+1)-1",
                      logit="exp(x)/(1+exp(x))",
                      identity="x")

obs2opts <- function(opts,obs) {opts$obs <- obs;opts} # Adds observed data to the model options.

p_transform_mat <- function(x) { # Transforms a matrix (typically representing variances and covariances) for optimization by converting it to a vector with appropriate transformations applied.
  ### Converts a matrix to a form that can be fed to optim
  ## matrix to list-format
  origx <- x
  if (is.matrix(x)) x <- mat_to_list(x)
  sames <- map_lgl(x,~all(. %in% c("same","Same","SAME")))
  if (length(sames)>0 && sames[1]) stop("Cannot start with the _same_ argument!")
  pn <- 0
  matmapping <- x
  for (i in seq_along(x)) {
    if (any(str_detect(x[[i]],"fix|Fix|FIX"))) {
      numbers <- parse_number(x[[i]])
      matmapping[[i]] <- paste0(numbers," fix")
      x[[i]] <- numbers
    } else if (!sames[i]) {
      pnseq <- pn+seq_along(x[[i]])
      pn <- tail(pnseq,1)
      matmapping[[i]] <- paste0("p",pnseq)
    } else {
      matmapping[[i]] <- matmapping[[i-1]]
      x[[i]] <- x[[i-1]]
    }
  }
  ## x2 <- list_to_mat(x)
  x2 <- list_to_mat(map(x,as.numeric))
  x3 <- cov2cor(x2);diag(x3) <- diag(x2) ## retain diagonals, offdiagonals to correlations
  matmapping <- list_to_mat(matmapping)
  parlocs <- which(grepl("p",c(matmapping)) & !duplicated(c(matmapping)))
  transformlist <- rep("log",length(parlocs))
  for (i in 2:length(transformlist)) {
    matind <- which(matmapping==paste0("p",i),arr.ind=TRUE)[1,]
    if (diff(matind)!=0) ## if non-diagonal
      transformlist[i] <- "covlogit"
  }
  values <- map_dbl(seq_along(parlocs),~do.call(transformlist[.],
                                                list(as.numeric(x3[parlocs[.]]))))
  backtransformfunc <- function(p) {
    if (missing(p)) p <- values
    res <- x2
    backfuns <- backtransfunchooser(transformlist)
    seqnow <- seq_along(p)
    pfill <- map_dbl(seqnow,~backfuns[[.]](p[.]))
    for (i in seqnow)
      res[paste0("p",i) ==matmapping] <- pfill[i]
    sdmat <- diag(sqrt(diag(res)))
    cormat <- res;diag(cormat) <- 1
    res2 <- sdmat %*% cormat %*% sdmat
    fixedvals <- grepl("fix",matmapping)
    res2[fixedvals] <- x2[fixedvals]
    res2
  }
  d_psi_d_psitrans_long <- function(p) {
    if (missing(p)) p <- values
    actvals <- backtransformfunc(p)
    ## then just return by map(p) actvals(p) for diagonals, and something a bit more elaborate for offdiags!
    res0 <- matrix(0,nrow(x2),ncol(x2))
    map(seq_along(p),function(i) {
      res <- res0
      indices <- which(matmapping==paste0("p",i),arr.ind=TRUE)
      if (transformlist[i]=="log") {
        res[indices] <- actvals[indices]
      } else {
        corr <- 2*exp(p[i])/(exp(p[i]) + 1)^2
        res[indices] <- corr*sqrt(actvals[indices[1,1],indices[1,1]]*actvals[indices[1,2],indices[1,2]])
      }
      res
    })
  }
  derivseq <- derivtransfunchooser(transformlist)
  d_psi_d_psitrans_short <- function(p) {
    if (missing(p)) p <- values
    d_psi_d_psitrans_long(p) %>% map(~unique(.[.!=0]))
  }
  d_bi_d_omega <- function(biseqt,pt) {
    if (missing(pt)) pt <- values
    m <- backtransformfunc(pt)
    map(seq_along(values),function(i) {
      pind <- paste0("p",i)
      if (pind %in% diag(matmapping)) {
        x <- rep(0,nrow(m))
        x[diag(matmapping)==pind] <- m[matmapping==pind]
        ifelse(x==0,0,biseqt * 1/sqrt(x)/2)
      } else {
        gen_bi_here <- function(p) {
          Omega <- m
          Omega[matmapping==pind] <- Omega[matmapping==pind]+p
          if (any(diag(Omega)==0))
            diag(Omega)[diag(Omega)==0] <- 1e-10
          biseqt %*% chol(Omega)
        }
        c(jacobiann(gen_bi_here,0))
      }
    })
  }
  mm <- matrix(0,nrow(x2),ncol(x2))
  d_omega_d_Omega <- map(paste0("p",seq_along(values)),function(i) {
    mm[matmapping==i] <- 1
    mm})
  list(values=values,backtransformfunc=backtransformfunc,transfunclist=transformlist,mapping=matmapping,
       d_psi_d_psitrans_long=d_psi_d_psitrans_long,d_psi_d_psitrans_short=d_psi_d_psitrans_short,
       d_bi_d_omega=d_bi_d_omega,d_omega_d_Omega=d_omega_d_Omega)
}

p_transform_vec <- function(x) { # Transforms a parameter vector for optimization, handling transformations like log, covlogit, and fixed parameters.
  ### Transforms a vector of input values into something that can be fed to optim
  ## params with "fix" are removed
  origx <- x
  fixed <- str_detect(x,"fix") | str_detect(x,"Fix") | str_detect(x,"FIX")
  x <- x[!fixed]
  ## if other strings are present, then they are assumed to relate to the desired transformation
  xtransfuns <- rep("log",length(x))
  othertransfuns <- str_extract(format(x,scientific=FALSE),"[[:alpha:]]+")
  xtransfuns[!is.na(othertransfuns)] <- na.omit(othertransfuns)[1]
  x2 <- str_extract(x,".*[[:digit:]]+") %>% as.numeric()
  values <- map_dbl(seq_along(x2),~do.call(xtransfuns[.],list(x2[.])))
  backtransseq <- backtransfunchooser(xtransfuns)
  derivseq <- derivtransfunchooser(xtransfuns)
  backtransformfunc <- function(p,numeric=TRUE) {
    if (missing(p)) p <- values
    p2 <- map_dbl(seq_along(p),~backtransseq[[.]](p[.]))
    res <- origx
    res[!fixed] <- p2
    if (numeric & !is.numeric(res)) as.numeric(str_extract(res,"[[:digit:] [:punct:]]+")) else res
  }
  d_psi_d_psitrans_long <- function(p) {
    if (missing(p)) p <- values
    res <- rep(0,length(origx))
    map(seq_along(p),function(i) {
      ii <- which(!fixed)[i]
      res[ii] <- derivseq[[i]](p[i])
      res
    })
  }
  d_psi_d_psitrans_short <- function(p) {
    if (missing(p)) p <- values
    map2(derivseq,p,~.x(.y)) %>% unlist()
  }
  list(values=values,backtransformfunc=backtransformfunc,transfunclist=xtransfuns,origx=origx,
       d_psi_d_psitrans_long=d_psi_d_psitrans_long,d_psi_d_psitrans_short=d_psi_d_psitrans_short)
}


p2opts <- function(opts,pp) {
  if (!is.list(pp)) {
    opts$pt <- pp
    opts$p <- opts$ptrans(pp)
  } else {
    opts$p <- pp
    opts$pt <- p_to_optim(pp)$values
  }
  return(opts)
}

samplogdensfun <- function(bi,p,omega_expansion) { # Computes the log-density of random effects bi under an expanded and unexpanded covariance matrix.
  map_dbl(1:nrow(bi),function(i) {
    proposal <- dmnorm(bi[i,],mean=rep(0,nrow(p$Omega)),sigma=p$Omega*omega_expansion,log=TRUE)$den
    true <- dmnorm(bi[i,],mean=rep(0,nrow(p$Omega)),sigma=p$Omega,log=TRUE)$den
    true-proposal
  })
}

upd_opts <- function(opts0,optslist) { # Initializes and generates core options and settings for modeling and optimization, including random effects, simulation settings, and likelihood approximations.
  ## note: this function always resets biseq and adist, and takes away obs!!
  if (!all(names(optslist) %in% names(formals(genopts))))
    stop("Argument not understood by genopts() function!")
  update_biseq <- TRUE
  if ("biseq" %in% names(optslist)) update_biseq <- FALSE
  for (i in names(optslist)) opts0[[i]] <- optslist[[i]]
  if (update_biseq) opts0$biseq <- NA;
  opts0$adist <- NULL
  opts0$ptrans <- NULL;opts0$pderiv <- NULL;
  opts0$ai <- NULL;opts0$pt <- NULL
  opts0$obs <- NULL
  opts0$d_g_d_beta <- NULL
  opts0$d_g_d_bi <- NULL
  opts0$d_bi_d_omega <- NULL
  opts0$d_omega_d_Omega <- NULL
  do.call(genopts,opts0)
}

uptolow <- function(m) { # Copies the upper triangular part of a matrix to the lower triangular and vice versa.
  m[lower.tri(m)] <- t(m)[lower.tri(m)]
  m
}

calculate_bsv <- function(back_transformed_params) {
  tryCatch({
    # Extract diagonal elements from the Omega matrix
    omega_diag <- diag(back_transformed_params$Omega)

    # Calculate BSV (CV%) for (log-)normal parameters
    #bsv_cv <- sqrt(exp(omega_diag) - 1) * 100
    bsv_cv <- sqrt(omega_diag) * 100

    # Handle potential NA cases or missing values in Omega
    c(bsv_cv, rep(NA, length(omega_diag) - length(bsv_cv)))
  }, error = function(e) {
    # In case of any error, return NA for all
    rep(NA, length(diag(back_transformed_params$Omega)))
  })
}

perturb_init <- function(init, pertubation) {
  init + rnorm(length(init), mean = 0, sd = pertubation * abs(init))  # Slight perturbation
}

compute_nll <- function(opts, params, nomap) {
  if (nomap) {
    maxfunc(p2opts(opts, params))(params)
  } else {
    Reduce('+', map(opts, ~ maxfunc(p2opts(., params))(params)))
  }
}

run_chain <- function(chain, opts, obs, init, maxiter, phase_fractions, convcrit_nll,
                      max_worse_iterations, pertubation, nomap) {
  chain_start_time <- Sys.time()  # Start time for the chain
  chain_init <- if (chain == 1) init else perturb_init(init, pertubation)

  if (nomap) {
    opts <- opts %>% p2opts(chain_init) %>% obs2opts(obs)
  } else {
    opts <- map(seq_along(opts), function(i) opts[[i]] %>% p2opts(chain_init) %>% obs2opts(obs[[i]]))
  }

  res <- tibble(
    iter = 1,
    parameters = vector("list", maxiter),
    nll = NA_real_,
    approx_nll = NA_real_,
    iteration_time = 0
  )

  res$parameters[[1]] <- chain_init

  if (nomap) {
    res$nll[1] <- maxfunc(opts)(chain_init)
  } else {
    res$nll[1] <- Reduce('+', map(opts, ~ maxfunc(.)(chain_init)))
  }

  res$approx_nll[1] <- res$nll[1]

  if (chain == 1) {  # Print live output for single-chain mode
    cat(sprintf("Chain %d:\nIter | %-12s\n", chain, sprintf("NLL and Parameters (%d values)", length(init))))
    cat(sprintf("%s\n", strrep("-", 80)))
    cat(sprintf("%4d: %s\n", 1, paste(sprintf("%8.3f", c(res$nll[1], chain_init)), collapse = " ")))
  }

  phase_params <- list(
    list(bounds = 2, maxeval = 5000, phase_name = "Wide Search Phase"),
    list(bounds = 1, maxeval = 5000, phase_name = "Focussed Search Phase"),
    list(bounds = 0.5, maxeval = 5000, phase_name = "Fine-Tuning Phase"),
    list(bounds = 0.01, maxeval = 5000, phase_name = "Precision Phase")
  )

  if (length(phase_fractions) != length(phase_params)) {
    stop("Number of phase_fractions must match the number of phases.")
  }

  cumulative_iters <- cumsum(c(floor(phase_fractions[-length(phase_fractions)] * maxiter), maxiter))
  phase_start_iters <- c(2, head(cumulative_iters, -1) + 1)
  phase_end_iters <- cumulative_iters

  best_nll <- res$nll[1]
  best_params <- res$parameters[[1]]
  current_iter <- 2

  for (phase_idx in seq_along(phase_params)) {
    current_phase <- phase_params[[phase_idx]]

    if (chain == 1) {
      cat(sprintf("\n### %s ###\n", current_phase$phase_name))
    }

    # Call handle_phase
    phase_result <- handle_phase(
      current_phase = current_phase,
      opts = opts,
      chain_init = best_params,
      best_params = best_params,
      best_nll = best_nll,
      current_iter = current_iter,
      phase_end_iter = phase_end_iters[phase_idx],
      nomap = nomap,
      convcrit_nll = convcrit_nll,
      max_worse_iterations = max_worse_iterations,
      res = res,
      maxiter = maxiter,
      chain = chain  # Pass the chain index for printing control
    )

    # Update variables for the next phase
    best_params <- phase_result$best_params
    best_nll <- phase_result$best_nll
    res <- phase_result$res
    current_iter <- phase_result$current_iter

    if (phase_result$phase_converged) {
      cat(sprintf("Phase %s converged at iteration %d.\n", current_phase$phase_name, current_iter - 1))
    }
  }

  res <- res[!is.na(res$nll), ]
  chain_time <- as.numeric(difftime(Sys.time(), chain_start_time, units = "secs"))

  cat(sprintf("\nChain %d Complete: Final NLL = %.3f, Time Elapsed = %.2f seconds\n \n",
              chain, best_nll, chain_time))

  return(list(res = res, best_nll = best_nll, best_params = best_params, time = chain_time))
}

handle_phase <- function(current_phase, opts, chain_init, best_params, best_nll, current_iter, phase_end_iter,
                         nomap, convcrit_nll, max_worse_iterations, res, maxiter, chain) {
  phase_converged <- FALSE
  worse_counter <- 0

  for (i in current_iter:phase_end_iter) {
    if (current_iter > maxiter) {
      cat("Maximum iterations reached. Terminating optimization.\n")
      break
    }

    iter_start_time <- Sys.time()

    if (nomap) {
      ff <- maxfunc(p2opts(opts, chain_init))
    } else {
      ffs <- map(opts, ~ maxfunc(p2opts(., chain_init)))
      ff <- function(p) Reduce('+', map(ffs, ~ .(p)))
    }

    m0 <- nloptr::nloptr(
      x0 = chain_init,
      eval_f = function(params) ff(params),
      lb = chain_init - current_phase$bounds,
      ub = chain_init + current_phase$bounds,
      opts = list(
        algorithm = "NLOPT_LN_BOBYQA",
        ftol_rel = .Machine$double.eps,
        maxeval = current_phase$maxeval
      )
    )

    # Optimization step
    chain_init <- m0$solution
    res$parameters[[current_iter]] <- chain_init
    res$nll[current_iter] <- compute_nll(opts, chain_init, nomap)
    res$approx_nll[current_iter] <- m0$objective
    res$iteration_time[current_iter] <- as.numeric(difftime(Sys.time(), iter_start_time, units = "secs"))

    # Update best parameters if a better NLL is found
    if (res$nll[current_iter] < best_nll) {
      best_nll <- res$nll[current_iter]
      best_params <- chain_init
    }

    # Printing live updates for the chain
    if (chain == 1) {  # Only print for the first chain
      cat(sprintf("%4d: %s\n", current_iter, paste(sprintf("%8.3f", c(res$nll[current_iter], chain_init)), collapse = " ")))
    }

    # Convergence check
    if (abs(res$nll[current_iter] - res$approx_nll[current_iter]) < convcrit_nll) {
      phase_converged <- TRUE
      current_iter <- current_iter + 1
      break  # Skip the rest of this phase, continue to the next phase
    }

    # Early phase termination due to worse counter
    worse_counter <- if (res$nll[current_iter] > best_nll) worse_counter + 1 else 0
    if (worse_counter >= max_worse_iterations) {
      current_iter <- current_iter + 1
      break  # Skip the rest of this phase, continue to the next phase
    }

    current_iter <- current_iter + 1  # Increment after every iteration
  }

  return(list(
    best_params = best_params,
    best_nll = best_nll,
    res = res,
    phase_converged = phase_converged,
    current_iter = current_iter
  ))
}
```
