## Supplementary material for "Aggregate data modelling: a fast implementation for fitting pharmacometrics models to summary-level data in R": electronic supplement: README.txt

These are the files in this folder:

run.R is the run file and will run the entire simulation and visualization of this manuscript.

cpp.R contains the pharmacometrics C++ models that are used in the simulations.
internal_functions.R copies the internal functions of the admr R package, which are necessary in these simulations.
Simulation files contain the functions to run each simulation. To run the files separately all libraries, parameters and functions need to be loaded in the environment.
Visualization files contain the functions to create the figures from the manuscript and extra.  To run the files separately all libraries, parameters and functions need to be loaded in the environment.

Feel free to reach out for clarification to the corresponding author.
