## Supplementary material for "Aggregate data modelling: a fast implementation for fitting pharmacometrics models to summary-level data in R": electronic supplement: run.R

```
library(admr)
library(tidyverse)
library(nlmixr2)
library(ggplot2)
library(dplyr)
library(Rcpp)
library(mnormt)
library(patchwork)
source("scripts/internal_functions.R")
source("scripts/cpp.R")


# Run simulations
nsim <- 20

## Simulation 1
Cpp_predder <- function(xt,theta_i,dose) {
  if (!is.matrix(theta_i) || nrow(theta_i)==1) {
    if (missing(dose)) dose <- 100
    Cpp_twocomp(theta_i,xt,FALSE)*dose
  } else {
    if (missing(dose)) dose <- rep(100,nrow(theta_i))
    Cpp_twocomp2(theta_i[,1],theta_i[,2],theta_i[,3],
                 theta_i[,4],theta_i[,5],xt,FALSE)* (dose %*% t(rep(1,length(xt))))
  }
}

opts1a <- genopts(time=c(.1,.25,.5,1,2,3,5,8,12),
                  p=list(beta=c(cl = 5, v1 = 10, v2 = 30,q = 10, ka = 1),Omega=omegas(.09,0,5),Sigma_prop=0.01),
                  nsim=2000,n=2000,
                  adist=NULL, interact=TRUE,fo_appr=FALSE,biseq=NA,omega_expansion=1,
                  f=Cpp_predder)

#source("scripts/simulation1.R")
source("scripts/visualization1.R")

## Simulation 2
Cpp_predder <- function(xt,theta_i,dose) {
  if (!is.matrix(theta_i) || nrow(theta_i)==1) {
    if (missing(dose)) dose <- 100
    Cpp_twocomp(theta_i,xt,FALSE)*dose
  } else {
    if (missing(dose)) dose <- rep(100,nrow(theta_i))
    Cpp_twocomp2(theta_i[,1],theta_i[,2],theta_i[,3],
                 theta_i[,4],theta_i[,5],xt,FALSE)* (dose %*% t(rep(1,length(xt))))
  }
}

opts1a <- genopts(time=c(.1,.25,.5,1,2,3,5,8,12),
                  p=list(beta=c(cl = 5, v1 = 10, v2 = 30,q = 10, ka = 1),Omega=omegas(.09,0,5),Sigma_prop=0.01),
                  nsim=1000,n=1000,
                  adist=NULL, interact=TRUE,fo_appr=FALSE,biseq=NA,omega_expansion=1,
                  f=Cpp_predder)
opts1b <- upd_opts(opts1a,list(p=list(beta=c(5,10,30,10,1),Omega=omegas(.09,0,3),Sigma_prop=0.01),
                               g=function(beta,bi=rep(0,length(beta)),ai) {beta*exp(c(bi[1:2],0,0,bi[3]))},
                               single_betas=c(F,F,T,T,F),
                               p_thetai=function(p,origbeta,bi) {
                                 dmnorm(bi,mean=log(p$beta[c(1,2,5)]/origbeta[c(1,2,5)]),sigma=p$Omega,log=TRUE)$den
                               }))

#source("scripts/simulation2.R")
source("scripts/visualization2.R")

## Simulation 3
Cpp_predder_dt <- function(time, theta_i){
  if (!is.matrix(theta_i) || nrow(theta_i)==1) {
    Cpp_transit_dt(theta_i,time)
  } else {
    Cpp_transit2_dt(theta_i[,1],theta_i[,2],theta_i[,3],
                    theta_i[,4],theta_i[,5],theta_i[,6],theta_i[,7],time)
  }
}
opts1c <- genopts(time = c(seq(168, 176, 0.2), seq(177, 190, 1)),
                  p=list(beta=c(cl = 2, vc = 10, vp = 30, q = 10, ka = 2, mtt= 0.5, n = 5),Omega=omegas(0.09,0,7),Sigma_prop=0.01),
                  nsim=1000,n=1000,
                  f=Cpp_predder_dt,
                  omega_expansion = 1)
#source("scripts/simulation3.R")
source("scripts/visualization3.R")


## Joint visualization
# boxplot
parameter_plot_paper1 <- parameter_plot_paper1 + 
  labs(title = "Simulation 1") +
  theme(plot.title = element_text(size = 16, face = "bold", hjust = 0.5))

parameter_plot_paper2 <- parameter_plot_paper2 + 
  labs(title = "Simulation 2") +
  theme(plot.title = element_text(size = 16, face = "bold", hjust = 0.5))

parameter_plot_paper3 <- parameter_plot_paper3 + 
  labs(title = "Simulation 3") +
  theme(plot.title = element_text(size = 16, face = "bold", hjust = 0.5))

combined_panel <- (parameter_plot_paper1 / parameter_plot_paper2 / parameter_plot_paper3) +
  plot_layout(ncol = 1)

combined_panel
ggsave("figures/boxplot_combined.svg", plot = combined_panel, width=8, height=10)


# vpc
vpc_plot_paper1 <- vpc_plot1 + 
  labs(title = "Simulation 1") +
  theme(plot.title = element_text(size = 16, face = "bold", hjust = 0.5))

vpc_plot_paper2 <- vpc_plot2 +
  labs(title = "Simulation 2") +
  theme(plot.title = element_text(size = 16, face = "bold", hjust = 0.5))

vpc_plot_paper3 <- vpc_plot3 +
  labs(title = "Simulation 3") +
  theme(plot.title = element_text(size = 16, face = "bold", hjust = 0.5))

combined_vpc_panel <- (vpc_plot_paper1 / vpc_plot_paper2 / vpc_plot_paper3) +
  plot_layout(ncol = 3)

combined_vpc_panel
ggsave("figures/vpc_combined.svg", plot = combined_vpc_panel, width=15, height=4.1)
```
