## Supplementary material for "Aggregate data modelling: a fast implementation for fitting pharmacometrics models to summary-level data in R": electronic supplement: simulation1.R

```
set.seed(1)

##################################
## Auxiliary functions for case ##
##################################
fitboth <- function(opts, nsim = 1, seedd = 1) {
  set.seed(seedd)
  initlist <- map(seq_len(nsim), ~ opts$pt - rnorm(length(opts$pt), sd = 0.1))
  ptrue <- opts$pt
  ptrue_originalscale <- opts$p
  res_IR_MC <- vector("list", nsim)
  p_IR_MC <- vector("list", nsim)
  res_bobyqa <- vector("list", nsim)
  p_bobyqa <- vector("list", nsim)
  #seedlist <- sample(0:500, nsim, replace = TRUE)
  obs <- MCapprEV(opts)
  function(estIR_MC = TRUE, estbobyqa = TRUE) {
    if (estIR_MC) {
      for (i in which(map_lgl(res_IR_MC, is.null))) {
        message(paste0("IR-MC case ", i))
        #obs <- meancov(gendataset(opts, seed = seedlist[i]))
        opts.test <- upd_opts(opts, list(nsim = 5000, omega_expansion = 1))
        res_IR_MC[[i]] <<- timedIRMC(initlist[[i]], opts.test, obs, convcrit_nll = 0.0001) %>%
          mutate(initset = i, iter = 1:n(), Method = "IR-MC") %>%
          { 
            .$p_orig <- lapply(.$p, opts$ptrans)
            .
          }
        p_IR_MC[[i]] <<- map(res_IR_MC[[i]]$p, ~ c(opts$ptrans(.x))) 
      }
    }
    
    if (estbobyqa) {
      for (i in which(map_lgl(res_bobyqa, is.null))) {
        message(paste0("MC case ", i))
        #obs <- meancov(gendataset(opts, seed = seedlist[i]))
        opts.test <- upd_opts(opts, list(nsim = 5000, omega_expansion = 1))
        res_bobyqa[[i]] <<- mutate(timedbobyqa(initlist[[i]], opts.test, obs), initset = i, Method = "MC", appr_nll = NA) %>%
          { 
            .$p_orig <- lapply(.$p, opts$ptrans)
            .
          }
        p_bobyqa[[i]] <<- map(res_bobyqa[[i]]$p, ~ c(opts$ptrans(.x)))
      }
    }
    
    # Combine results and calculate differences for every iteration
    do.call(rbind, c(res_IR_MC, res_bobyqa)) %>%
      rowwise() %>%
      mutate(
        pdiff = list(p - ptrue),
        pdiff_originalscale = list(
          if (Method == "IR-MC") {
            map2(p_IR_MC[[initset]][[iter]], ptrue_originalscale, ~ .x - .y)
          } else {
            map2(p_bobyqa[[initset]][[iter]], ptrue_originalscale, ~ .x - .y)
          }
        )
      )
  }
}

#######################################
## Speedup of the IR-MC algorithm
#######################################
## Define the model
f1a <- fitboth(opts1a,nsim)

res1a <- f1a() %>% mutate(Case = "A") %>%
  unnest_wider(pdiff_originalscale, names_sep = "_") %>%
  mutate(
    # Extract beta as a vector with names
    beta_vector = purrr::map(pdiff_originalscale_beta, ~{
      beta_values <- as.vector(.x)
      names(beta_values) <- c("beta1", "beta2", "beta3", "beta4", "beta5")
      beta_values
    }),
    
    # Extract Omega diagonal from 5x5 matrices with names
    Omega_diag_vector = purrr::map(pdiff_originalscale_Omega, ~{
      Omega_diag_values <- diag(.x)[1:5]
      names(Omega_diag_values) <- paste0("Omega", 1:5)
      Omega_diag_values
    }),  # Diagonal from 5x5 matrix
    
    # Extract Sigma_prop as a vector with names
    Sigma_prop_vector = purrr::map(pdiff_originalscale_Sigma_prop, ~{
      Sigma_prop_values <- as.vector(.x)
      names(Sigma_prop_values) <- "Sigma_prop"
      Sigma_prop_values
    })
  ) %>%
  # Combine all vectors into one vector for each row, keeping names
  mutate(
    pdiff_originalscale = purrr::map2(beta_vector, Omega_diag_vector, ~c(.x, .y)) %>%
      purrr::map2(Sigma_prop_vector, ~c(.x, .y)),
    
    # Calculate relative error for each parameter
    relative_error = purrr::map(pdiff_originalscale, ~{
      # Define the true values for each row
      true_beta <- c(cl = 5, v1 = 10, v2 = 30, q = 10, ka = 1)
      true_Omega <- c(Omega1 = 0.09, Omega2 = 0.09, Omega3 = 0.09, Omega4 = 0.09, Omega5 = 0.09)
      true_Sigma_prop <- 0.01
      
      # Calculate relative error for beta, Omega, and Sigma_prop
      beta_error <- .x[1:5] / true_beta * 100 
      Omega_error <- .x[6:10] / true_Omega * 100
      Sigma_prop_error <- .x[11] / true_Sigma_prop * 100
      
      # Combine the errors into one vector
      c(beta_error, Omega_error, Sigma_prop_error)
    })
  ) %>%
  # Keep only the relevant columns
  dplyr::select(p, nll, appr_nll, time, iter, initset, Method, pdiff, Case, pdiff_originalscale, relative_error)

# View the result with relative error
head(res1a)

saveRDS(res1a, "data/res1a.rds")
```
