## Supplementary material for "Aggregate data modelling: a fast implementation for fitting pharmacometrics models to summary-level data in R": electronic supplement: visualization3.R

```
res <- readRDS("data/res1c.rds")

# Define Leiden colors
leiden_colors <- c("#0c2577", "#4a7184", "#8592bb", "#dfe3ee", "#06123b")

bind_rows(filter(res,initset==1, Method=="IR-MC",),
          filter(res,initset==1, Method=="MC")) %>% 
  group_by(Method) %>% mutate(time=time-time[1]) %>% 
  unnest_longer(pdiff) %>% 
  ggplot(aes(unclass(time),exp(pdiff),col=pdiff_id,group=paste(Method,pdiff_id)))+
  geom_line()+scale_y_log10()+facet_grid(Method~.)+theme(legend.position="none")+
  xlab("Time (s)")+ylab("Ratio estimate/true")

res %>%
  group_by(Case,initset,Method) %>%
  summarize(time=tail(time,1)-head(time,1)) %>%
  spread(Method,time) %>% mutate(timeratio=unclass(MC)/unclass(`IR-MC`)) %>%
  group_by(Case) %>%
  summarize(meanMC=mean(MC),`meanIR-MC`=mean(`IR-MC`),meanratio=mean(timeratio),
            sdMC=sd(MC),`sdIR-MC`=sd(`IR-MC`),sdratio=sd(timeratio))


# Function to create convergence plots
convplots <- function(d) {
  d %>%
    unnest_longer(pdiff) %>%
    group_by(initset) %>%
    mutate(time = as.numeric(time - time[1])) %>%
    ggplot(aes(time, pdiff, group = factor(initset))) +
    geom_hline(yintercept = 0, col = 2) +
    geom_line() +
    facet_wrap(~pdiff_id)
}

# Filter data for bobyqa and IR-MC
MC_data <- filter(res, Case == "A", Method == "MC")
IR_MC_data <- filter(res, Case == "A", Method == "IR-MC")

convplots(MC_data)
convplots(IR_MC_data)


# Create a named vector to replace pdiff_id abbreviations
param_names <- c("cl" = "Clearance",
                 "v1" = "Volume Central",
                 "v2" = "Volume Peripheral",
                 "q" = "Intercompartmental Clearance",
                 "ka" = "Absorption Rate", 
                 "mtt" = "Mean Transit Time",
                 "n" = "Number of Transit Compartments",
                 "Omega1" = "Clearance Omega",
                 "Omega2" = "Volume Central Omega",
                 "Omega3" = "Volume Peripheral Omega",
                 "Omega4" = "Intercompartmental Clearance Omega",
                 "Omega5" = "Absorption Rate Omega", 
                 "Omega6" = "Mean Transit Time Omega",
                 "Omega7" = "Number of Transit Compartments Omega",
                 "Sigma_prop" = "Sigma Proportional Error")

final_plot_paper <- res %>%
  unnest_longer(pdiff, indices_to = "pdiff_id") %>%
  filter(pdiff_id %in% c("ka", "Omega5", "Sigma_prop")) %>%
  mutate(
    Method = factor(Method, levels = c("MC", "IR-MC")), # Ensure MC is first
    pdiff_id = recode(pdiff_id, !!!param_names) # Replace abbreviations with descriptive names
  ) %>%
  group_by(Method, initset) %>% # Group by Method and initset to calculate time offsets
  mutate(
    time = as.numeric(time - min(time)) # Align time so each Method starts at 0
  ) %>%
  ggplot(aes(time, pdiff, group = interaction(initset, Method), color = Method)) +
  geom_hline(yintercept = 0, col = leiden_colors[5]) +
  geom_line() +
  facet_grid(Method ~ pdiff_id, scales = "fixed") + # Method in rows, pdiff_id in columns
  theme_minimal() +
  theme_bw(base_size=16) +
  scale_color_manual(values = c(leiden_colors[1], leiden_colors[2])) + # Use first two colors for MC and IR-MC
  labs(x = "Time (s)", y = "Parameter Difference") +
  theme(
    legend.position = "none", # Remove legend
    panel.border = element_rect(color = "black", fill = NA, size = 0.5), # Add boxes around plots
    strip.background = element_rect(fill = leiden_colors[4], color = "black") # Background color for facet labels
  )

final_plot_appendix1 <- res %>%
  unnest_longer(pdiff, indices_to = "pdiff_id") %>%
  filter(pdiff_id %in% c("cl", "v1", "v2", "q", "ka", "mtt", "n", "Sigma_prop")) %>%
  mutate(
    Method = factor(Method, levels = c("MC", "IR-MC")), # Ensure MC is first
    pdiff_id = recode(pdiff_id, !!!param_names) # Replace abbreviations with descriptive names
  ) %>%
  group_by(Method, initset) %>% # Group by Method and initset to calculate time offsets
  mutate(
    time = as.numeric(time - min(time)) # Align time so each Method starts at 0
  ) %>%
  ggplot(aes(time, pdiff, group = interaction(initset, Method), color = Method)) +
  geom_hline(yintercept = 0, col = leiden_colors[5]) +
  geom_line() +
  facet_grid(Method ~ pdiff_id, scales = "fixed") + # Method in rows, pdiff_id in columns
  theme_minimal() +
  theme_bw(base_size=16) +
  scale_color_manual(values = c(leiden_colors[1], leiden_colors[2])) + # Use first two colors for MC and IR-MC
  labs(x = "Time (s)", y = "Parameter Difference") +
  theme(
    legend.position = "none", # Remove legend
    panel.border = element_rect(color = "black", fill = NA, size = 0.5), # Add boxes around plots
    strip.background = element_rect(fill = leiden_colors[4], color = "black") # Background color for facet labels
  )

final_plot_appendix2 <- res %>%
  unnest_longer(pdiff, indices_to = "pdiff_id") %>%
  filter(pdiff_id %in% c("Omega1", "Omega2", "Omega3", "Omega4", "Omega5", "Omega6", "Omega7")) %>%
  mutate(
    Method = factor(Method, levels = c("MC", "IR-MC")), # Ensure MC is first
    pdiff_id = recode(pdiff_id, !!!param_names) # Replace abbreviations with descriptive names
  ) %>%
  group_by(Method, initset) %>% # Group by Method and initset to calculate time offsets
  mutate(
    time = as.numeric(time - min(time)) # Align time so each Method starts at 0
  ) %>%
  ggplot(aes(time, pdiff, group = interaction(initset, Method), color = Method)) +
  geom_hline(yintercept = 0, col = leiden_colors[5]) +
  geom_line() +
  facet_grid(Method ~ pdiff_id, scales = "fixed") + # Method in rows, pdiff_id in columns
  theme_minimal() +
  theme_bw(base_size=16) +
  scale_color_manual(values = c(leiden_colors[1], leiden_colors[2])) + # Use first two colors for MC and IR-MC
  labs(x = "Time (s)", y = "Parameter Difference") +
  theme(
    legend.position = "none", # Remove legend
    panel.border = element_rect(color = "black", fill = NA, size = 0.5), # Add boxes around plots
    strip.background = element_rect(fill = leiden_colors[4], color = "black") # Background color for facet labels
  )

final_plot_paper
final_plot_appendix1
final_plot_appendix2

speed_boxplot <- res %>%
  group_by(Case, initset, Method) %>%
  summarize(time = tail(time, 1) - head(time, 1)) %>%
  mutate(Method = factor(Method, levels = c("IR-MC", "MC"))) %>%  # Ensure IR-MC is first
  ggplot(aes(x = time, y = Method, fill = Method)) +  # Map Method to y-axis
  geom_boxplot(alpha = 0.7) +  # Boxplot by Method
  scale_fill_manual(values = c(leiden_colors[2], leiden_colors[1])) +  # Reverse Leiden colors
  labs(x = "Time (s)", y = "") +  # Remove y-axis label
  theme_minimal() +
  theme_bw(base_size=16) +
  theme(
    legend.position = "none",  # Remove legend
    panel.border = element_rect(color = "black", fill = NA, size = 0.5),  # Box around plot
    strip.background = element_rect(fill = leiden_colors[4], color = "black")  # Facet label background
  )

# Display the boxplot
print(speed_boxplot)


# Create a named vector to replace pdiff_id abbreviations
param_names <- c("beta1" = "Clearance",
                 "beta2" = "Volume Central",
                 "beta3" = "Volume Peripheral",
                 "beta4" = "Intercompartmental Clearance",
                 "beta5" = "Absorption Rate", 
                 "beta6" = "Mean Transit Time",
                 "beta7" = "Number of Transit Compartments",
                 "Omega1" = "Clearance Omega",
                 "Omega2" = "Volume Central Omega",
                 "Omega3" = "Volume Peripheral Omega",
                 "Omega4" = "Intercompartmental Clearance Omega",
                 "Omega5" = "Absorption Rate Omega", 
                 "Omega6" = "Mean Transit Time Omega",
                 "Omega7" = "Number of Transit Compartments Omega",
                 "Sigma_prop" = "Sigma Proportional Error")


parameter_plot_paper3 <- res %>%
  unnest_longer(relative_error, indices_to = "pdiff_id") %>%
  filter(pdiff_id %in% c("beta5", "Omega5", "Sigma_prop")) %>%
  mutate(
    Method = factor(Method, levels = c("MC", "IR-MC")), # Ensure MC is first
    pdiff_id = recode(pdiff_id, !!!param_names) # Replace abbreviations with descriptive names
  ) %>%
  group_by(Method, initset, pdiff_id) %>%
  filter(nll == min(nll)) %>%  # Select only the last iteration for each combination
  summarize(avg_pdiff = mean(relative_error, na.rm = TRUE)) %>%  # Calculate average pdiff for last iteration
  ggplot(aes(x = Method, y = avg_pdiff, fill = Method)) +  # Boxplot by Method
  geom_boxplot(alpha = 0.7, color = "black", position = position_dodge(width = 0.75)) +  # Side-by-side boxplots for each Method
  geom_hline(yintercept = 0, linetype = "dashed", color = "red", size = 0.8) +  # Horizontal line at y = 1
  scale_fill_manual(values = c(leiden_colors[1], leiden_colors[2])) +  # Leiden colors for MC and IR-MC
  facet_wrap(~pdiff_id, scales = "fixed") +  # Facet by each parameter (ka, Omega5, Sigma_prop)
  labs(x = "Method", y = "Relative bias (%)") +  # Axis labels
  theme_minimal() +
  theme_bw(base_size=16) +
  theme(
    legend.position = "none",  # Remove legend
    panel.border = element_rect(color = "black", fill = NA, size = 0.5),  # Box around plot
    strip.background = element_rect(fill = leiden_colors[4], color = "black")  # Facet label background
  ) +
  coord_cartesian(ylim = c(-5, 5))  # Set y-axis limits from 0.99 to 1.01


parameter_plot_appendix1 <- res %>%
  unnest_longer(relative_error, indices_to = "pdiff_id") %>%
  filter(pdiff_id %in% c("beta1", "beta2", "beta3", "beta4", "beta5", "beta6", "beta7", "Sigma_prop")) %>%
  mutate(
    Method = factor(Method, levels = c("MC", "IR-MC")), # Ensure MC is first
    pdiff_id = recode(pdiff_id, !!!param_names) # Replace abbreviations with descriptive names
  ) %>%
  group_by(Method, initset, pdiff_id) %>%
  filter(nll == min(nll)) %>%  # Select only the last iteration for each combination
  summarize(avg_pdiff = mean(relative_error, na.rm = TRUE)) %>%  # Calculate average pdiff for last iteration
  ggplot(aes(x = Method, y = avg_pdiff, fill = Method)) +  # Boxplot by Method
  geom_boxplot(alpha = 0.7, color = "black", position = position_dodge(width = 0.75)) +  # Side-by-side boxplots for each Method
  geom_hline(yintercept = 0, linetype = "dashed", color = "red", size = 0.8) +  # Horizontal line at y = 1
  scale_fill_manual(values = c(leiden_colors[1], leiden_colors[2])) +  # Leiden colors for MC and IR-MC
  facet_wrap(~pdiff_id, scales = "fixed") +  # Facet by each parameter (cl, v1, v2, q, ka, Sigma_prop)
  labs(x = "Method", y = "Relative bias (%)") +  # Axis labels
  theme_minimal() +
  theme(
    legend.position = "none",  # Remove legend
    panel.border = element_rect(color = "black", fill = NA, size = 0.5),  # Box around plot
    strip.background = element_rect(fill = leiden_colors[4], color = "black"),  # Facet label background
    axis.title = element_text(size = 16),
    axis.text = element_text(size = 14),
    strip.text = element_text(size = 14)
  ) +
  coord_cartesian(ylim = c(-5, 5))  # Set y-axis limits from 0.99 to 1.01


parameter_plot_appendix2 <- res %>%
  unnest_longer(relative_error, indices_to = "pdiff_id") %>%
  filter(pdiff_id %in% c("Omega1", "Omega2", "Omega3", "Omega4", "Omega5", "Omega6", "Omega7")) %>%
  mutate(
    Method = factor(Method, levels = c("MC", "IR-MC")), # Ensure MC is first
    pdiff_id = recode(pdiff_id, !!!param_names) # Replace abbreviations with descriptive names
  ) %>%
  group_by(Method, initset, pdiff_id) %>%
  filter(nll == min(nll)) %>%  # Select only the last iteration for each combination
  summarize(avg_pdiff = mean(relative_error, na.rm = TRUE)) %>%  # Calculate average pdiff for last iteration
  ggplot(aes(x = Method, y = avg_pdiff, fill = Method)) +  # Boxplot by Method
  geom_boxplot(alpha = 0.7, color = "black", position = position_dodge(width = 0.75)) +  # Side-by-side boxplots for each Method
  geom_hline(yintercept = 1, linetype = "dashed", color = "red", size = 0.8) +  # Horizontal line at y = 1
  scale_fill_manual(values = c(leiden_colors[1], leiden_colors[2])) +  # Leiden colors for MC and IR-MC
  facet_wrap(~pdiff_id, scales = "fixed") +  # Facet by each parameter (Omega1, Omega2, Omega3, Omega4, Omega5)
  labs(x = "Method", y = "Relative bias (%)") +  # Axis labels
  theme_minimal() +
  theme(
    legend.position = "none",  # Remove legend
    panel.border = element_rect(color = "black", fill = NA, size = 0.5),  # Box around plot
    strip.background = element_rect(fill = leiden_colors[4], color = "black"),  # Facet label background
    axis.title = element_text(size = 16),
    axis.text = element_text(size = 14),
    strip.text = element_text(size = 14)
  ) +
  coord_cartesian(ylim = c(-10, 10))  # Set y-axis limits from 0.99 to 1.01


parameter_plot_paper3
parameter_plot_appendix1
parameter_plot_appendix2


ggsave("figures/boxplot3.svg", plot = parameter_plot_paper3, width=7.2, height=4)


## VPC PLOTS
vpcdata <- res %>%
  filter(Method == "IR-MC") %>%
  mutate(
    Method = factor(Method, levels = c("MC", "IR-MC"))
  ) %>%
  group_by(initset, Method) %>%
  filter(nll == min(nll)) %>%
  ungroup() 

# 1. Set up simulation
time_points <- seq(168, 190, by = 0.1)  # Dense time points for smooth curves
n_subjects <- 10000  # Number of subjects to simulate

# True parameters
true_beta <- c(cl = 2, v1 = 10, v2 = 30, q = 10, ka = 2, mtt= 0.5, n = 5)
true_omega <- diag(rep(0.09, 7))


# Generate random effects for true model
mv <- MASS::mvrnorm(n_subjects, rep(0, 7), true_omega)

# Create parameter matrix for true model
theta_true <- matrix(nrow = n_subjects, ncol = 7)
for(i in 1:n_subjects) {
  theta_true[i,] <- c(
    true_beta["cl"] * exp(mv[i,1]),
    true_beta["v1"] * exp(mv[i,2]),
    true_beta["v2"] * exp(mv[i,3]),
    true_beta["q"]  * exp(mv[i,4]),
    true_beta["ka"] * exp(mv[i,5]),
    true_beta["mtt"] * exp(mv[i,6]),
    true_beta["n"] * exp(mv[i,7])
  )
}

# Simulate true model using Cpp_predder
true_predictions <- Cpp_predder_dt(time_points, theta_true)

# Convert to data frame and calculate percentiles for true model
true_predictions_df <- data.frame(
  time = rep(time_points, each = n_subjects),
  concentration = as.vector(true_predictions),
  sim.id = rep(1:n_subjects, times = length(time_points))
)

true_percentiles <- true_predictions_df %>%
  group_by(time) %>%
  summarize(
    median = median(concentration),
    lower = quantile(concentration, 0.05),
    upper = quantile(concentration, 0.95)
  ) %>%
  ungroup()

# Function to simulate from IR-MC parameters
simulate_irmc <- function(params, n_subjects) {
  # Extract fixed effects and variance components
  beta <- params$beta
  omega <- params$Omega
  
  # Generate random effects
  mv <- MASS::mvrnorm(n_subjects, rep(0, 7), omega)
  
  # Create parameter matrix for all subjects
  theta_i <- matrix(nrow = n_subjects, ncol = 7)
  for(i in 1:n_subjects) {
    theta_i[i,] <- c(
      beta[1] * exp(mv[i,1]),
      beta[2] * exp(mv[i,2]),
      beta[3] * exp(mv[i,3]),
      beta[4] * exp(mv[i,4]),
      beta[5] * exp(mv[i,5]),
      beta[6] * exp(mv[i,6]),
      beta[7] * exp(mv[i,7])
    )
  }
  
  # Simulate concentrations using Cpp_predder
  pred <- Cpp_predder_dt(time_points, theta_i)
  return(pred)
}


# Generate predictions for IR-MC iterations
irmc_predictions <- vpcdata %>%
  ungroup() %>%  # First ungroup the data
  as.data.frame() %>%  # Convert to regular data frame
  dplyr::select(initset, p) %>%  # Use dplyr::select explicitly
  rowwise() %>%
  mutate(
    params = list(opts1c$ptrans(p)),  # Transform parameters
    predictions = list(simulate_irmc(params, n_subjects))
  ) %>%
  ungroup() %>%
  mutate(
    data = map2(predictions, initset, ~{
      data.frame(
        time = rep(time_points, each = n_subjects),
        concentration = as.vector(.x),
        sim.id = rep(1:n_subjects, times = length(time_points))
      )
    })
  ) %>%
  dplyr::select(initset, data) %>%  # Use dplyr::select explicitly
  unnest(data)

# Calculate percentiles for IR-MC iterations
irmc_percentiles <- irmc_predictions %>%
  group_by(initset, time) %>%
  summarize(
    median = median(concentration),
    lower = quantile(concentration, 0.05),
    upper = quantile(concentration, 0.95)
  ) %>%
  ungroup()

observed_times <- c(seq(168, 175, 0.2), seq(176, 190, 1))
observed_data <- true_predictions_df %>%
  filter(time %in% observed_times) %>%
  group_by(time) %>%
  summarize(
    concentration = median(concentration)
  ) %>%
  ungroup()


# Add 'source' and dummy 'initset' to match columns
irmc_percentiles$source <- "Iterations"
true_percentiles$source <- "True Model"
true_percentiles$initset <- NA  # To match irmc_percentiles

# Combine the data
combined_data <- rbind(
  irmc_percentiles[, c("time", "median", "lower", "upper", "initset", "source")],
  true_percentiles[, c("time", "median", "lower", "upper", "initset", "source")]
)


# Add source labels
irmc_percentiles$source <- "Iterations"
true_percentiles$source <- "True model"

# Add dummy initset if needed
if (!"initset" %in% names(true_percentiles)) {
  true_percentiles$initset <- NA
}


vpc_plot3 <- ggplot() +
  # IR-MC iterations lines (no legend)
  geom_line(data = irmc_percentiles, 
            aes(x = time, y = median, color = "Aggregate model"), size = 1) +
  geom_line(data = irmc_percentiles, 
            aes(x = time, y = lower, color = "Aggregate model"), size = 1) +
  geom_line(data = irmc_percentiles, 
            aes(x = time, y = upper, color = "Aggregate model"), size = 1) +
  
  # Observed data points with color mapping for legend
  geom_point(data = observed_data, 
             aes(x = time, y = concentration, color = "Observed timepoints"),
             size = 2, shape = 15) +
  
  # True model lines with color mapping for legend
  geom_line(data = true_percentiles, 
            aes(x = time, y = median, color = "True model"), size = 1, linetype = "dashed") +
  geom_line(data = true_percentiles, 
            aes(x = time, y = lower, color = "True model"), size = 1, linetype = "dashed") +
  geom_line(data = true_percentiles, 
            aes(x = time, y = upper, color = "True model"), size = 1, linetype = "dashed") +
  
  labs(
    x = "Time (hours)",
    y = "Plasma concentration (mg/L)",
    color = NULL
  ) +
  scale_color_manual(values = c("True model" = "#f46e32", "Aggregate model" = leiden_colors[1], "Observed timepoints" = "black")) +
  theme_bw(base_size=16) +
  theme(
    panel.border = element_rect(color = "black", fill = NA, size = 0.5),
    panel.grid.major = element_blank(),
    panel.grid.minor = element_blank()
  ) +
  scale_x_continuous(breaks = seq(168, 190, by = 4)) +
  scale_y_continuous(trans = "log10", limits = c(0.1, 8))

print(vpc_plot3)

ggsave("figures/vpcplot3.svg", plot = vpc_plot3, width=9.2, height=5.6)
```
